# Multi-Task Batteries for Precision Functional Mapping

**DOI:** 10.64898/2026.03.20.713227

**Authors:** Bassel Arafat, Caroline Nettekoven, Jinkang Derrick Xiang, Jörn Diedrichsen

**Author notes:** Corresponding author: Jörn Diedrichsen.

## Abstract

Functional brain mapping is an important tool to understand the organization of the human brain, both at the group level, but also to an increasing degree at the level of the individual. There are currently two main approaches to do so. Resting-state fMRI relies on inter-regional correlations of random fluctuations of the signal. In contrast, task-based localizers typically use a single-contrast between a task of interest and a matched control task to identify the location of a functional region in an individual brain. In this paper, we propose and evaluate a third approach: the use of multi-task batteries for both localization of a single functional region and parcellation of multiple functional regions. We show that multi-task localizers produce more consistent estimation of a single functional region across subjects than the single-contrast approach using the same amount of fMRI data. Furthermore, we demonstrate that the multi-task approach is sensitive to true inter-individual differences in region size, and does not suffer the same influence of signal-to-noise ratio that biases the single-contrast localizer. We then address the question of how to select tasks for the battery, and present a data-driven strategy that optimizes the characterization of a brain structure of interest. We show that such batteries outperform randomly selected batteries both for building individual parcellations as well as individual connectivity models. Finally, we demonstrate that an interspersed design - where all tasks are presented in each imaging run - yields more reliable results than splitting the tasks across different runs. We present an open source toolbox for the implementation of multi-task batteries, along with a library containing group-averaged activity patterns that can be used to optimize battery selection for different brain structures of interest.

## 1 INTRODUCTION

Functional magnetic resonance imaging (fMRI) is an important non-invasive tool to understand the organization of the human brain. A number of influential group-level atlases have been developed (Yeo et al., 2011; Glasser et al., 2016; Gordon et al., 2016; Schaefer et al., 2018), which have greatly advanced our understanding of the brain’s broad organization. However, such group-based maps ignore the substantial inter-individual variability in the finer aspects of this organization (Mueller et al., 2013; Laumann et al., 2015; Gordon et al., 2017; Gratton et al., 2018; Braga and Buckner, 2017). This limitation has sparked a growing interest in the field for individualized brain mapping, which aims to characterize functional brain regions in single subjects. Such precision mapping is important when studying specific functional regions, but also for clinical applications, such as surgical planning and identifying individualized treatments.

One common approach to individualized brain mapping is resting-state fMRI (rsfMRI), where subjects lie in the scanner while spontaneous blood-oxygen-level-dependent (BOLD) signal fluctuations are recorded. Based on the correlations of these fluctuations, one can group different brain locations into functional networks. This approach can also be used to characterize brain organization at the level of the individual (Gordon et al., 2017; Wang et al., 2015; Kong et al., 2019, 2021; Xue et al., 2021). rsfMRI is popular due to its ease of implementation and applicability across a broad range of healthy and clinical populations. One key limitation of this method, however, is that the inter-regional correlations do not only reflect neural interactions but also non-neural noise sources such as head motion, respiration, heart rate and scanning artifacts (Buckner et al., 2013; Reid et al., 2019; Chen et al., 2025; Biswal and Uddin, 2025), which can bias the results.

A second approach is to use task-based fMRI, in which the BOLD signal is measured while participants perform one or more tasks. Alternating between tasks in a randomized design allows experimenters greater control over measurement noise. Traditionally, task-based fMRI for brain mapping has been applied using a single-contrast approach. Here, a single functional region of interest (ROI) is identified by comparing an experimental and a control task to target a specific function of interest (Kanwisher et al., 1997; Dodell-Feder et al., 2011; Scott et al., 2017). For example, to isolate the language network, reading sentences is contrasted with reading non-words (Fedorenko et al., 2010). This approach, however, can only serve to identify a single brain region, and does not offer the same comprehensive parcellation of individual brains that can be derived from rsfMRI.

As an alternative to these two approaches, we propose to use a wide battery of tasks to localize functional regions in individual brains (Barch et al., 2013; Pinho et al., 2018; King et al., 2019). In this multi-task approach, a battery with a diverse set of tasks is designed to sample a broad range of brain functions. In a companion paper (Nettekoven et al., 2026), we demonstrate that brain parcellations derived from such multi-task data outperform brain parcellations derived from resting-state data in predicting the brain activity patterns of novel tasks, even when the two approaches use an equivalent amount of data from the same participants. This result can be demonstrated to generalize across a range of brain structures, model types and datasets. Importantly, we also show that the result depends strongly on the exact choice of tasks, if the battery only consists of a few tasks. With an increasing number and diversity of tasks, however, the functional parcellation becomes increasingly independent of the specific tasks chosen and can be used to find a general task-invariant parcellation of the brain.

In this paper, we will further evaluate the power of the multi-task approach and address a number of important decisions when designing a multi-task battery. In the first part of the paper we show that a multi-task battery approach has distinct advantages over the single-contrast localizer, even if we only want to identify a single functional ROI. Specifically, we show that the multi-task approach allows for a better estimation of individual region sizes, which can be an important metric for describing inter-individual differences.

In the second part of the paper, we will then address the question of how many and which tasks should ideally be selected for a multi-task battery. While this decision will often be guided by specific functional and practical considerations, we propose a complementary, data-driven approach. Using a library of candidate tasks, and their associated brain activity maps, we can ask which strategy of task selection would lead to an optimal characterization of a specific brain structure. This approach is especially useful if the scanning time and the number of tasks for each individual are restricted for practical reasons. We evaluate different ways of choosing a task battery in the context of two practical applications of brain mapping: Subdividing a brain structure comprehensively into a set of functionally distinct regions (brain parcellation; Nettekoven et al., 2024; Zhi et al., 2025), and identifying the connections between two different brain structures (connectivity models; King et al., 2023; Cole et al., 2016; Reid et al., 2019).

In the final part of the paper, we then consider different experimental design choices of how the chosen tasks should be implemented during an fMRI scanning session. We compare a design in which similar tasks are grouped into separate runs (Barch et al., 2013), to one in which all tasks are randomly interspersed within every imaging run (King et al., 2019). We also consider order and carry-over effects, both of which have consequences for the exact design choices.

To make multi-task batteries easy to build and run, we are presenting a Python-based toolbox that implements more than 20 tasks and that allows for the flexible and fast assembly of new batteries. The toolbox is accompanied by a library of group-averaged activity patterns, enabling data-driven optimal battery selection for any brain structure of interest.

## 2 METHODS

### 2.1 Datasets: Language Task Battery

To compare the traditional functional localization approach to multi-task functional localization, we designed a multi-task battery that focused on language-related tasks. The dataset included 16 tasks (including rest), some of which contained multiple conditions, yielding a total of 18 task conditions (see supplementary table S2 for task descriptions). Within the task battery, we included traditional language localizer tasks: sentence reading, non-word reading, intact passage listening and degraded passage listening (Fedorenko et al., 2010; Scott et al., 2017). We collected data for 17 participants (9 female, 8 male, mean age = 22.5, SD = 3.1) across 8 imaging runs. In each imaging run, each task was performed once in random order. Each task started with a 5s instruction period, in which the name of the task and the response assignment to the buttons was shown on the screen. This was followed by 30 seconds of performing the task. The hand used for responses was alternated across runs, with the exception of the finger-sequence task, which required responses from both hands in all runs. The index and middle fingers of both hands were placed on the button pressing box throughout the whole experiment.

Each participant underwent a behavioral training session 1-7 days before their fMRI scanning session. Each training session started with verbally familiarizing the participants with the requirements of each task. This was followed by a training period where participants trained each task that required explicit responses (finger sequence, n-back, demand grid and oddball) separately. After the training period, each participant performed 3 training runs with all tasks, each equivalent to an imaging run. Mean accuracy across participants averaged across the 3 training runs was 91.14% (±1.22% SEM). Mean accuracy during the scanning session was 85.64% (±1.21% SEM). This training program ensured that participants were familiar with the requirements for each task and had experience in performing them before the fMRI scanning session. For both the training session and the scanning session, participants were provided trial-specific feedback (green fixation cross for correct and red fixation cross for incorrect) for tasks where feedback can be given (see supplementary table S2 for task feedback). At the end of each run, participants were presented with a scoreboard that showed their overall accuracy and reaction time for tasks that require a button press response.

fMRI data were collected on a 7T Siemens scanner at the Center for Functional and Metabolic Mapping at Western University. For the anatomical image, we acquired a high-resolution magnetization prepared rapid acquisition gradient echo (MP2RAGE) scan with an isotropic voxel size of 0.7mm. For functional images, we used echo-planar imaging with a multi-band factor 2 and in-plane acceleration of 3. Whole brain data were acquired covering both the neocortex and the cerebellum. Imaging parameters were: TR = 1.1s, TE = 20 ms, slices = 56, voxel size = 2.3mm isotropic. Each run had a length of 523 TRs (575s). For four subjects, scanning parameters were slightly different (TR = 1.3, volumes = 443, slices = 64).

### 2.2 Datasets: The Multi-Domain Task Battery (MDTB)

To evaluate different task selection strategies, we used the Multi-Domain Task Battery (MDTB) dataset (King et al., 2019). The dataset included two task sets (A and B). Task Set A included 17 tasks (29 task conditions) and task set B included 17 tasks (32 task conditions). 14 task conditions were shared between the two sets. Twenty-four participants (16 female, 8 male, mean age = 23.8, SD = 2.6) were scanned during 4 sessions, two sessions for each task set. Each session consisted of 8 10min runs. Within each run, the 17 tasks were presented in random order, starting with a 5-second instruction period and 30 seconds of performing the task. Images were acquired on a Siemens 3T MRI scanner with a voxel resolution of 2.5 × 2.5 × 3.0 mm^3^ and a TR of 1s (see King et al. (2019) for a detailed description of the dataset and tasks).

### 2.3 Datasets: The Human Connectome Project (HCP)

To compare different experimental designs, we also used the task-based fMRI data from the Human Connectome Project (HCP) (Barch et al., 2013). We used the minimally preprocessed 3T task-fMRI data from the HCP 1200 subjects release (2017), selecting 50 of the 100 unrelated-subjects group (25 female, 25 male, mean age estimated based on age ranges = 29.7, SD = 3.7). This dataset included two imaging runs for each of seven task sessions (working memory, gambling, motor, language, social, relational and emotion). For details, see Barch et al. (2013).

### 2.4 fMRI processing and estimation of activity profiles

For the Language and Multi-domain task battery datasets, data were preprocessed using SPM12 and custom MATLAB code separately for each dataset. Functional data were realigned for head motion. The mean functional image was then co-registered to the anatomical image and this transformation was applied to all functional volumes. No smoothing or group normalization was applied at this stage. A general linear model (GLM) was fitted to the time series of each voxel for each imaging run. The 5 second instruction periods were modeled using a single regressor per run. Each task was then modeled using a boxcar function of 30s or multiple regressors if a block contained multiple task conditions. The 30s rest block was explicitly modeled as a task. Each regressor was then convolved with the standard hemodynamic response function.

For the HCP dataset we used the minimally preprocessed version of the data (Glasser et al., 2013). A GLM was fitted to the time series of each voxel for each imaging run and session using the FEAT design files provided with the HCP 1200 subjects data 2017 release. No spatial smoothing or normalization was applied at this point.

For group analysis, the data were brought into *SUIT* space (for the cerebellum) and into *fs32k* space (for the neocortex). For *SUIT* normalization, the cerebellum and brainstem were isolated from the remainder of the brain and then normalized to the cerebellum-only SUIT template (Diedrichsen, 2006). For *fs32k*, the individual neocortical surfaces were reconstructed using freesurfer (Fischl, 2012), and aligned to the spherical surface template based on sulcus depth and curvature information (Glasser et al., 2013). Beta weights from the first-level GLM were univariately pre-whitened by dividing each voxel’s estimate by the square root of the residual mean-square image. We used these pre-whitened betas as measures of task activation for all analyses in this paper. For all three datasets, task activation was expressed relative to the resting baseline in the same run. We then averaged the activity values across all available runs to obtain one activation value per task. Each voxel’s activation profile was then a vector containing the activation values for all unique tasks.

We also analyzed the activity profiles at the parcel level, using the parcels from two group atlases: the Glasser neocortical atlas (Glasser et al., 2016), which contains 180 parcels per hemisphere, and the NettekovenSym-32 cerebellar atlas (Nettekoven et al., 2024), which contains 16 parcels per hemisphere. Voxel-wise activity profiles were then averaged within each parcel across voxels.

### 2.5 Functional signal to noise ratio

To simulate functional data with realistic characteristics, we defined the functional signal-to-noise-ratio (fSNR) of individual activity estimates (per imaging run based on 30s of task data) as follows: We modeled the estimated activity value for each task condition *i*, voxel *p*, and imaging run *j*, as composed of a true activity value (the signal, *u_i,p_*) and run-specific measurement noise (*ε_i,p,j_*):

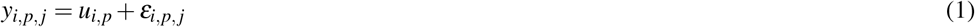

The true activity patterns have a variance of 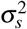across tasks and voxels, whereas the noise has a variance of 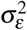across runs. We then define the fSNR of individual activity estimates as

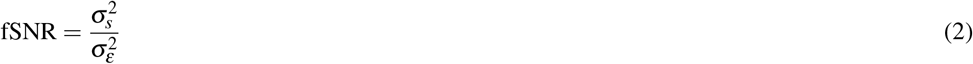

On real data, the size of these two variances can be estimated from the average variances and covariances between activity patterns. The average covariance of the estimates of the same tasks across two different runs serves as an estimator of the signal variance. The average variance of the activity patterns across all runs, tasks and voxels is the sum of the signal and noise variances. The noise variance can therefore be estimated by subtracting the signal variance from this quantity.

We estimated the fSNR for each subject in the MDTB dataset by extracting the estimated neocortical activity patterns for each task, run and vertex in *fs32k* space. For the language dataset we extracted the activity patterns for each task, run, and voxel in the cerebellum in *SUIT* space. We then used the covariance matrix of activity estimates across runs to estimate the signal and noise variances and then calculated the fSNR for each individual.

To capture the typical distribution of fSNRs, a gamma distribution was fitted to the estimated fSNR values. For the MDTB dataset, the best-fitting distribution had the parameters *α* = 6.77, and *β* = 0.0027.

### 2.6 Simulations: general methods

To simulate the data we would acquire for each hypothetical task set or functional localizer under realistic measurement conditions, we assumed that we had 8 min of individual scan time. We then allocated an equal amount of time for each unique task. We assume that if we allocated 90s to a task, we would have 3 repeated task phases of 30s and would average across these estimates. As the measurement variance for the average reduces proportionally with the number of independent samples, we can predict the effective fSNR for the average estimate based on *t* seconds from the fSNR of the 30s estimates (see section 2.5 for details on fSNR estimation):

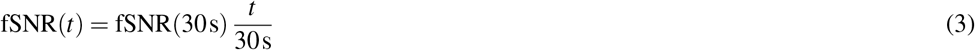

For the parcellation and functional localization simulations, we generate data on a 30×30 pixel grid, with each voxel assigned to one of five functionally distinct regions. The region assignment was encoded in the region ×voxel matrix **U**, with **U***_k,i_* = 1 if the *i^th^* voxel was assigned to the *k^th^* parcel.

Each region was then associated with a true functional profile across *N* tasks, **v***_k_*. Each of these vectors had a squared norm of *N*. The functional profiles of the different regions were then concatenated into the *N* ×*K* matrix **V**. We then generated artificial activity data **Y** by adding normally distributed noise (*ε*) across each task and voxel with mean 0 and a variance of 1*/f SNR*:

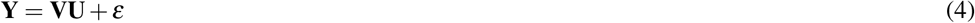

For connectivity simulations, we generated the source structure data only using a set of simulated activity profiles for 5 hypothetical brain locations across tasks **V**. To generate the target structure data **Y**, we randomly sampled a connectivity weights matrix **W** that relates each of the 5 source structure regions to 100 target structure regions. Noise *ε* was also sampled from a normal distribution with a mean of 0 and variance set to 1*/f SNR*:

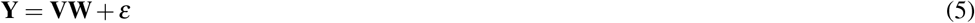

### 2.7 Functional localization: Simulations

To compare single-contrast and multi-task approaches to functional localization under controlled conditions, we simulated data across five regions (see 2.6 for details), with one region serving as the target region to be localized. 1000 individual-specific parcellations **U***_i_* were generated by introducing random variation in the size of the target region through adding or removing boundary pixels, yielding target region sizes uniformly distributed between 120 and 240 pixels with an average size of 180 pixels.

We then randomly generated the functional profile **V** for these 5 regions across 5 tasks. The norm of these functional profiles was 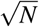, and they were orthogonal across the different regions. To achieve this we generated a 5 ×5 matrix of independent normal random variables and orthogonalized it using QR decomposition.

We implemented 3 types of localizers: (1) a single-contrast localizer with a fixed absolute threshold (“contrast-fixed”), (2) a single-contrast localizer with an adaptive percentile threshold (“contrast-adaptive”), and (3) a multi-task localizer that assigns each pixel to the region with the most similar multi-task response profile. For the single-contrast localizers, we chose the two tasks that maximized the contrast between the target region and the average of all the other regions. For the multi-task localizer, we used all five tasks. To capture the inter-individual differences in fSNR, we sampled the individual fSNR values from the fitted distribution to the estimated empirical fSNR values of the MDTB dataset (see 2.5). Using these values, we then simulated activity patterns of 8 min length for each battery for each individual (see 2.6 for details).

Based on simulated data, we then estimated the target region for each individual subject. For the multi-task localizer, each pixel was assigned to the parcel whose multi-task activation profile was most similar to the pixel’s activation profile, as measured by cosine similarity between profiles. For the single-contrast localizers, the target region was determined as all voxels exceeding a specific threshold on the single-contrast. The threshold was calibrated such that the average estimated target region size matched the average true region size across individuals. Localization accuracy was quantified using the Dice coefficient (Dice, 1945) between the true target region and the estimated ROI for each individual.

### 2.8 Functional localization: empirical example

To compare single-contrast and multi-task localizers, we used the language task battery dataset. For the single-contrast localizer, we computed the contrast between sentence reading and non-word reading for each subject, using the unsmoothed activity estimates from all 8 runs (8 minutes of imaging data, 4 minutes per task). We conducted a paired t-test for this contrast across runs, yielding a single t-value map per individual. These maps were thresholded across a range of t-values (0.1-2.8). At the upper limit of this range, fewer than 100 voxels remained active across subjects.

To assess the relationship between ROI size and data quality, we computed subject-specific fSNR values from the cerebellar data (see 2.5), excluding the sentence reading and non-word reading tasks. For each threshold we quantified the number of cerebellar voxels exceeding the threshold in each individual and correlated these values with the subject-specific fSNR estimates.

For the direct comparison to a multi-task localizer, we used the same dataset to assemble a battery consisting of five tasks: sentence reading, theory of mind, tongue movement, demand grid and action observation (action) (see supplementary table S2 for task descriptions). These tasks were chosen based on previous group functional atlases showing that they activate language regions and adjacent areas (Nettekoven et al., 2024). The amount of scanning data included was equivalent to 8 minutes of data (1.6 min per task).

We then parcellated the right cerebellar hemisphere of each individual into 5 regions, according to their functional profiles. To define these regions we used the right hemisphere of the Nettekoven-Sym32 group atlas (Nettekoven et al., 2024) but combined the 16 fine-grained regions for the right hemisphere into 5 coarser subregions, reflecting the major functional domains. We combined all motor regions (M1-M4), all action-observation regions (A1-A3) and all demand regions (D1-D4) into single motor, action and demand regions. We grouped sociolinguistic regions S1 and S2 into a single language region, as both show higher activity for sentence reading relative to non-word reading. The remaining sociolinguistic regions (S3-S5) defined the last and fifth parcel.

Based on the 5 region coarse atlas we estimated the functional profiles for these regions (see 2.4). Voxel and region activity profiles were mean centered and normalized across the five chosen tasks. We used a leave-one-subject-out cross-validation procedure, where the functional profiles were estimated using all subjects except one, and the voxels for that individual were then assigned to the parcel whose activity profile showed the highest cosine similarity with the voxel’s own activity profile.

### 2.9 Battery selection: Task library

In the second part of the paper, we tested different data-driven approaches to optimally design a task battery by choosing tasks from a larger library. In general, we consider a library of *L* tasks, each associated with an activity pattern measured across *K* brain regions, denoted **V***_L_*. All of these activity patterns are measured relative to a resting baseline.

For our simulations we generated a random library of 100 tasks that had random activation values across 5 regions, drawn from an independent standard normal distribution. For the real data examples, we used the measured activity pattern from 32 task conditions of task set B from the MDTB dataset, extracted from the brain structure of interest (neocortex or cerebellum). All activity patterns were expressed relative to the resting baseline.

To construct a task battery of *B* tasks, we simply select *B*− 1 rows of **V***_L_*. One of the rows of the task battery was always rest, and the activity values of this task were set to zero. In this way we constructed batteries of size *B* ranging from 3 to 16 unique tasks.

### 2.10 Battery selection: selection strategies

We aimed to compare two different criteria of selecting tasks for a battery based on the task activity patterns within the brain structure of interest. The first criterion *Maximal contrast* was designed to maximize the functional contrasts between the selected tasks and the resting baseline. Mathematically, we selected the task battery that maximized the variance of the functional contrasts across regions and tasks:

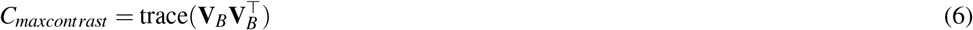

The second criterion *Minimal collinearity* was designed to select tasks that activated the target structure in maximally different ways. To achieve this, the variance of each activity pattern against the mean of the other tasks needed to be large, and the covariances between these activity patterns across regions needed to be low. Mathematically this can be achieved by first subtracting the mean activity from each brain location (the column mean of **V**_*B*_), resulting in the centered battery matrix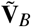. Note that the activity for rest vs. the mean of the other tasks is included in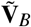. If we used the columns of this matrix as independent variables in a multiple regression analysis (as we do in the connectivity analysis), we know that the variance of the estimated parameter scales with the trace of the inverse gram matrix. To have all criteria be maximal for the best battery, we use here the negative trace:

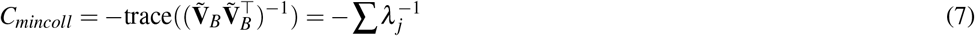

Where *λ_j_* are the eigen-values of 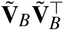. If 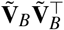 was not invertible, we used only the non-zero eigen-values of that matrix. As can be seen from Eq. 6, 7, both criteria depend on the task-by-task matrix 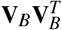, in one case without subtracting the mean, in the other case with subtracting the mean.

For real data, there are two ways of estimating this matrix: First, we can use the group-averaged activity patterns for each brain structure as rows in **V**_*B*_ and compute 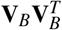 directly from that. Second, if all the activity patterns in the Library are measured in the same set of individuals, we can estimate the task-by-task matrix using the run-wise activity patterns from each individual participant:

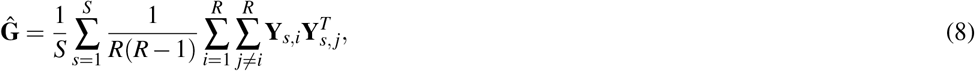

where **Y***_s,i_* is the measured activity in subject *s* in run *i* for the task (rows) and brain locations of interest (columns). This estimate differs from the simpler group-activity estimate in two ways: First, it captures differences between tasks that are only visible at the individual subject level, but which are obscured at the group-level, because two functional regions can be tightly interdigitated in a variable way across individuals (Braga and Buckner, 2017; Xue et al., 2021). Second, the estimate is cross-validated (by multiplying activity estimates from different runs only), such that measurement noise does not inflate the sums-of-squares of each individual contrast (the diagonal of **Ĝ**). While we used the second approach for greater precision in this paper, we found that the two approaches yield very similar results (see supplementary 7.1). This allows us to build a task-library by combining tasks that are measured in separate groups of individuals.

Finding the best battery from a larger library of tasks, however, is computationally prohibitive, as there are too many possible combinations for an exhaustive search. We therefore randomly built 10000 candidate batteries, and found the best battery among these according to each selection strategy. These batteries were then compared to those obtained via a random selection strategy, in which we sampled tasks without replacement at random without consideration of the activity patterns. This allowed us to assess the benefit of data-driven selection strategies over a naïve baseline in downstream individual brain mapping applications.

### 2.11 Battery selection: Simulations

To assess how each battery selection strategy influenced the quality of subsequent analyses, we conducted simulations estimating either a parcellation or a connectivity model from synthetic task battery data. Each simulation started by generating a random task-library of 100 tasks (see section 2.9) and five regions. We then chose the best task battery of a specific size given one of the two criteria, or selected a task battery at random.

For the **parcellation** simulations, we generated synthetic data for each battery **Y**_*b*_ on a 30 ×30 voxel grid (see section 2.6), where each voxel was assigned to one of five regions. Here, we only simulated a single individual. To estimate functional parcellations for each task battery we used the known activity profiles **V**_*b*_ to assign each voxel to the region that matched the measured activity profile best (i.e. has the highest cosine similarity). Both **V**_*b*_ and 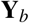 were mean-centered and normalized. To evaluate parcellation accuracy, we computed the Dice overlap between the estimated parcellation for the battery and the ground-truth parcellation, averaged across all 5 regions.

For the **connectivity** simulations, we used the true activity profile of the 5 regions as our source structure, generated random connectivity weights and from that the measured activities in the target structure (see section 2.6). From the generated data, we then estimated the connectivity weights **Ŵ**, using ridge regression. Source structure regressors were standardized to unit variance before fitting. We used a fixed regularization parameter *α* = 1. We evaluated the estimated connectivity model by calculating the Pearson correlation between the vectorized estimated connectivity weights **Ŵ**and the ground-truth connectivity weights **W**. This correlation reflects how well the task battery enables recovery of the true connectivity weights between a source and a target structure.

### 2.12 Battery selection: empirical application

To validate our battery selection simulations with real fMRI data, we applied the same framework to the MDTB dataset. As task set B contains a larger number of unique task conditions (32), we used it as the task library from which to construct candidate batteries. To control for the amount of total scan time, all batteries were constructed using 8 min of imaging data. This resulted in an artificial dataset for each task battery and subject for the brain structure of interest 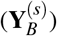. Based on these data, we then estimated the individual parcellations and connectivity models. We evaluated the quality of the estimated models by determining how well they could predict the activity pattern on task set A, which contained 14 shared and 15 unique task conditions.

In the **parcellation** analysis, we obtained individual parcellations within either the neocortex or the cerebellum. We first calculated the average functional profiles **V**_*B*_ within each region using either the Glasser atlas for neocortical parcellations or the NettekovenSym-32 atlas for cerebellar parcellations (see section 2.4) from all individuals, except the one we were parcellating. Based on the individual task battery data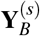, we then computed the cosine similarity between each voxel /vertex and the functional profile of each parcel in **V**_*B*_. We assigned each voxel/vertex to the parcel most similar in functional profile. The columns of **V**_*B*_ and 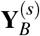were mean-centered and normalized before estimating parcellations.

We evaluated the quality of each individual parcellation by determining how well we could predict independent test data from the same individual 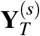. To derive predictions, we used the individual parcellation obtained from task set B, combined with the functional profile of each of the regions for task set A, estimated on all other subjects using a leave-one-subject-out procedure. The functional profiles were estimated from the group parcellation (Glasser or NettekovenSym-32, see section 2.4). The predicted activity profile for each voxel /vertex was then simply the functional profile of the region that brain location was assigned to in the individual parcellation. As an evaluation criterion, we computed the cosine similarity between the true and predicted test data for each subject and voxel and then averaged across voxels within each subject.

In the **connectivity** analysis, we estimated a directed connectivity model between the neocortex and the cerebellum (King et al., 2023) using each task battery. We then evaluated how well the connectivity model was able to predict task activity patterns in the cerebellum based on neocortical activity patterns on a different set of tasks (task set A). For estimating the connectivity model, the neocortical data of each individual *s* were summarized using a regular parcellation into a total of 1002 icosahedral regions, yielding a data matrix 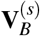. The cerebellar data were extracted in SUIT space, yielding a data matrix 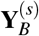. We estimated subject-specific connectivity weights, **Ŵ** ^(*s*)^, using ridge regression with a fixed regularization parameter *α* = *e*^8^, chosen based on prior work (King et al., 2023).

To evaluate the quality of the models, we used the estimated connectivity weights for each subject **Ŵ** ^(*s*)^together with the neocortical activity patterns from task set A 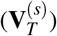to predict cerebellar activity patterns for the same tasks 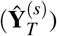. Model performance was measured as the cosine similarity between predicted and observed activity patterns. Cosine similarity was computed for each voxel and averaged across voxels within each subject.

### 2.13 Experimental design: Estimating variance caused by baseline estimation

As a typical example of a multi-task dataset with grouped design, we used the HCP task dataset (Barch et al., 2013). In this dataset, we determined how much run-to-run variability in activity estimates for each task is caused by the estimation of baseline. For this, we used the activity pattern of all tasks from the two available imaging runs for each session. All activity values were relative to the implicit baseline (rest). For each participant and session, we extracted the unsmoothed neocortical activity patterns in fs32k space, and computed the covariance matrix of these estimates across all vertices. The expected covariance between the activity pattern measured for the *i^th^* task in run 1 (*y*_*i*,1_) and in run 2 (*y_i_*,_2_) is determined only by the variance (strength) of the true signal pattern for that task.

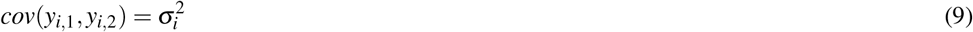

The expected covariance between the activity patterns between two different tasks (*i* and *j*) across runs is determined by the similarity of the true signal patterns.

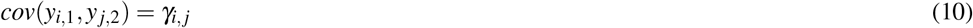

Within each run, the expected covariance between two tasks is again determined by the similarity of the two tasks. However, because the two tasks are estimated relative to the same baseline, the expected covariance is increased by the variability of that baseline estimate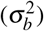:

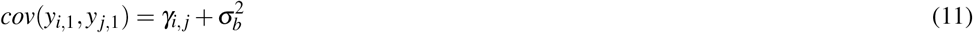

Finally, the within-run variance of each activity estimate is determined by the variance of the true signal pattern for task *i*, the baseline variance, and the variance of the noise unique to that run and task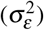:

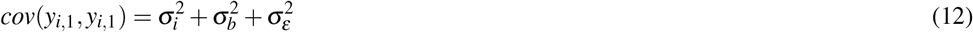

The covariances were computed from the estimated activity patterns across all vertices and then averaged across tasks. From the four different types of covariances, we used Eq. 9-12 to estimate the underlying variances for each participant and session.

### 2.14 Experimental design: Grouped vs. interspersed

Based on these estimates, we then compared two experimental designs for a multi-task dataset. In the **grouped** design, each imaging run contains a subset of tasks, typically from the same domain. By using different runs for different subsets of tasks, all tasks are covered. In the **interspersed** design, all tasks are repeated within each imaging run. Both designs also contain periods of rest within each run.

We then conducted simulations to compare the standard deviation of the contrast estimates of the two designs. Both designs had four unique tasks and rest, presented across two functional runs of 300s each. The difference between the two designs lies in whether tasks are grouped into separate runs or all tasks are included in each run.

Design matrices were created for each design while systematically varying the proportion of time points allocated to rest within each run. Each design matrix **X** contained a column for each task and imaging run plus an intercept regressor for each run. Assuming independent and identically distributed measurement noise across fMRI images with variance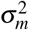, the variance-covariance matrix of the resultant beta-estimates (Worsley and Friston, 1995) is given by:

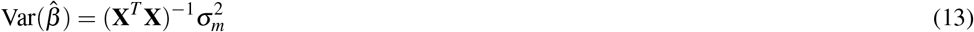

We evaluated the standard deviation of three types of contrasts: (i) the contrast between two tasks, conducted within each imaging run (for the interspersed design, the contrast was averaged across the two imaging runs); (ii) the contrast between two tasks, conducted across two imaging runs (for the grouped design only); and (iii) the contrast between each task and the implicit resting baseline. Each contrast is a linear combination of activity estimates and can therefore be expressed using a contrast vector **c**. The variance of each contrast can therefore be calculated as:

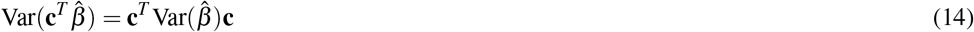

The standard deviation for each type of contrast was then calculated as the square root of the variance of each type of contrast.

To address the potential drawback of the interspersed design requiring an additional instruction period for each task, we conducted an additional simulation that allocated 5s of scan time to the instruction period for each 30s task block in the interspersed design (See S3). The grouped design remained unchanged.

### 2.15 Experimental design: Temporal autocorrelation

To estimate the influence of low-frequency fMRI fluctuations on the activity estimates in multi-task designs, we used Task Set A from the MDTB dataset, which consisted of 16 runs of 10 min length, each containing the same 17 tasks in different orders. We first expressed the activity patterns relative to the resting baseline (one of the 17 tasks) and then subtracted the mean pattern for each task (averaged across runs), thereby obtaining estimates of the run-wise residuals. These residuals were then sorted by the order in which they were presented in each run. We then calculated the covariance between all pairs of residuals, starting from pairs that immediately followed each other (task position difference = 1) up to residuals that were maximally separated (task position difference = 16). As the task signal is subtracted out (Eq. 9, 10), the covariance between tasks within-run is driven only by the variability of the common baseline estimate across runs.

### 2.16 Experimental design: Estimating variance caused by task carry-over

In interspersed multi-task designs, a potential concern is that the estimated activity for a given task might include residual activation from the preceding task. To estimate the magnitude of this carry-over effect, we used session A of the MDTB dataset. We used the covariance of the activity estimates for each task across runs to estimate the variance (strength) of the true signal pattern for that task (Eq. 9). This analysis was performed using the neocortical data in fs32k space. Because the order of tasks was randomized across runs (rather than counterbalanced for sequential effects), 5.2 % of these task activity pairs had the same preceding task. The difference between the covariance between task estimates with the same previous task and the covariance between task estimates with different previous tasks can therefore be taken as an estimate of the variance induced by a systematic carry-over effect.

We first conducted the analysis within each individual, thereby identifying carry-over effects that occurred in that individual. To determine what part of these carry-over effects were systematic across individuals, we then repeated the analysis, this time calculating the covariances for pairs of activity patterns from different individuals, and then averaging over all possible pairs of individuals. This allowed us to quantify the true signal variance of the carry-over effects in relation to the true signal variance of the task effect for both the group and within individuals.

## 3 RESULTS

### 3.1 Single-contrast vs. multi-task localizers: Simulation

The traditional approach to functional localization is to use a contrast between a task that contains the function of interest (Fig. 1a, Task a) and a control task (Task b) (Kanwisher et al., 1997). For example, to isolate the language network, reading sentences is contrasted with reading non-words (Fedorenko et al., 2010). To decide which voxels are considered part of the region of interest, the researcher needs to define an activity threshold, and all the voxels exceeding this threshold are assigned to the target region (Fig. 1a).

**Figure 1.**
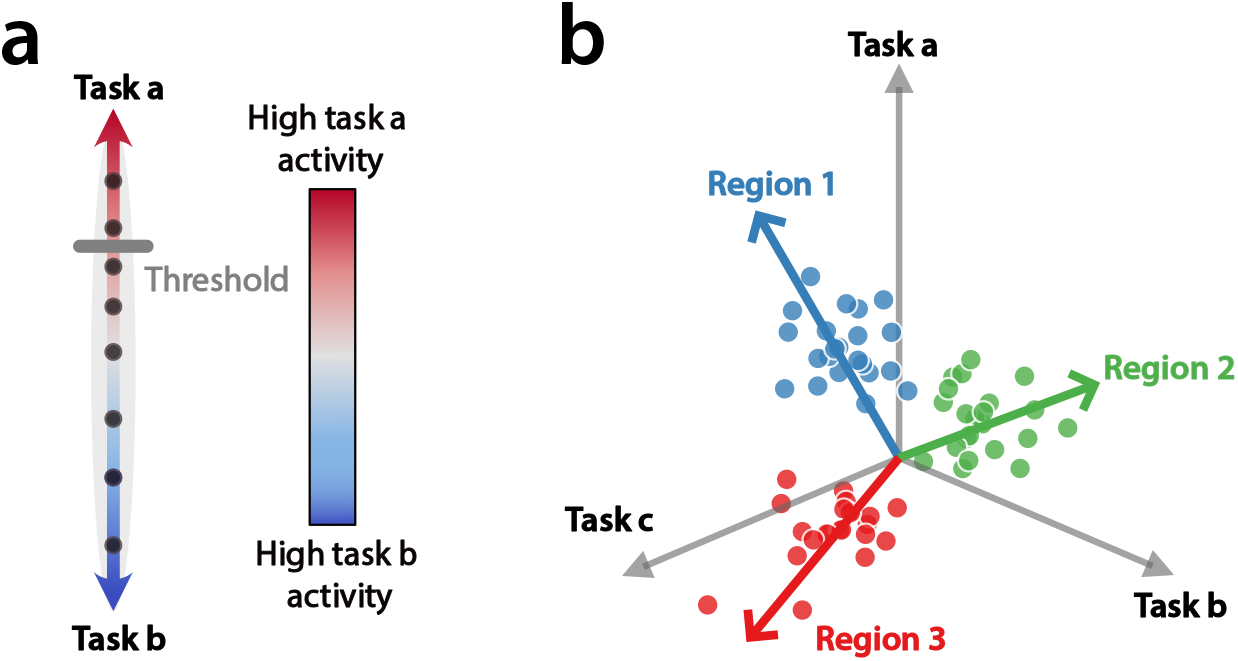
Single-contrast vs multi-task functional localization. **(a)** Single-contrast localizer. The activity of each voxel (dot) is characterized along a single dimension which compares its activity for Task a relative to a control task (Task b). All voxels exceeding a certain activity threshold (gray line) are selected as the desired ROI. **(b)** Multi-task localizer. The activity of each voxel is characterized by its response to multiple tasks relative to the mean across all tasks. Voxels located at the origin of the coordinate system respond equally to Task a, b and c. Different regions are defined by their response profile across all tasks (red, green and blue arrows). Voxels are assigned to the region with the most similar profile, independent of the length of their activity profile (distance from origin).

One option for choosing this threshold is to use a fixed statistical value. This approach, however, has the problem that the estimated size of the selected region will depend on the functional signal-to-noise ratio (fSNR, see 2.5) - for subjects with higher fSNRs, more voxels will meet the statistical threshold and the selected region will therefore be larger. In real datasets, fSNR varies by a factor of 3-4 across different subjects (Fig. 2a), leading to substantial differences in the size of the selected region. To remedy this situation, researchers often use an adaptive threshold, for example selecting a specific percentage of the highest activated voxels in each individual. This ensures that the size of the selected region will be the same across individuals (relative to the overall size of the brain structure). Of course, this approach will be insensitive to true inter-individual differences in the size of the functional region.

**Figure 2.**
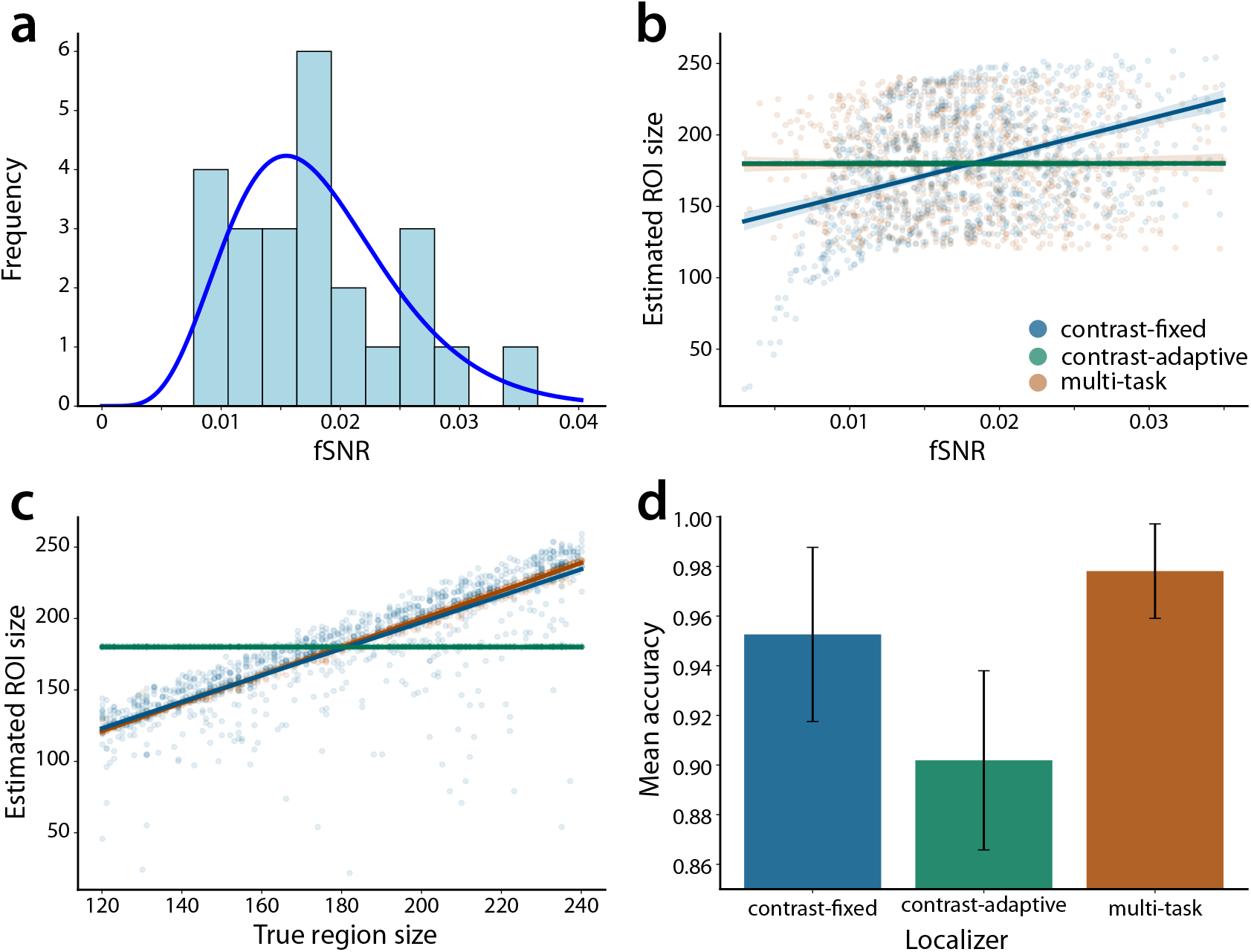
Simulated comparison of localization methods. **(a)** Distribution of subject-specific fSNRs estimated from the MDTB dataset. The line indicates the best-fitting gamma distribution. **(b)** Estimated ROI size as a function of fSNR for the three localizers. Each dot represents one simulated subject. Regression lines are shown for each of the localizers. **(c)** Estimated ROI size as a function of true ROI size for the three localizers. Each dot represents one simulated subject. Regression lines are shown for each of the localizers. **(d)** Average localization accuracy across localizer methods. Accuracy measured as the Dice coefficient between the estimated ROI and true region across simulated individuals. Error bars indicate across-subjects standard deviation of the differences between localizer methods.

To address these issues, multi-task localizers characterize each voxel according to its response to multiple tasks, with resting baseline phases treated as an independent task condition. The response profile (with the mean across tasks removed) then determines the location of each voxel in a multi-dimensional space (Fig. 1b). A voxel at the origin of the space would show the same value for all tasks (including rest), i.e. it would show no response. The distance from the origin indicates the strength of the response, and the direction of the vector characterizes the response profile across tasks. In this framework, each region can be characterized by a typical response profile (arrows). Instead of thresholding, each voxel is assigned to the single parcel with the most similar response profile (i.e. the one with the smallest angle). This has the advantage that the voxel assignment is not sensitive to the fSNR, which only influences the distance of the voxel from the origin, but not its direction.

To illustrate the difference between the two approaches, we simulated functional data across many individuals for a hypothetical brain structure, in which we aim to identify a target region surrounded by four other functional regions. While the general spatial layout was simulated to be similar across individuals, the size of the target region varied by a factor of two. Each region was also associated with a typical response profile across 5 hypothetical tasks. We then simulated functional data using fSNRs sampled from a realistic distribution (Fig. 2a, see 2.7).

We then tested the three approaches to functional localization. For the single-contrast localizer we used either a fixed statistical threshold (contrast-fixed), or an adaptive threshold defined by a specific percentage of pixels in each individual (contrast-adaptive). For the single-contrast localizers, the two tasks were selected to maximize the contrast between the target region and the average response of all the other regions. The threshold used for both single-contrast types was calibrated such that, on average, the estimated ROI size matched the true size of the target region across individuals (∼180 pixels). For the multi-task localizer all 5 tasks were used. The length of the simulated fMRI data was the same for the single-contrast and multi-task localizers - i.e. for the latter we used less data per task than for the former. For the contrast-fixed localizer, we found that the estimated ROI size varied systematically with fSNR (Fig. 2b). Individuals with higher fSNR values showed larger estimated ROIs, reflected in a positive correlation between fSNR and estimated ROI size (*r* = 0.41). In contrast, for the multi-task localizer, ROI size was independent of fSNR (*r* = 0.004), indicating its robustness to different fSNRs across individuals. For the contrast-adaptive localizer, the estimated ROI size did not depend on fSNR since it was a fixed number of pixels across all individuals. Consequently, this approach overestimated small regions and underestimated large regions, making it insensitive to true inter-individual differences in region size (Fig. 2c). Across all individuals, the multi-task localizer showed the highest accuracy when compared to both single-contrast approaches (Fig.2d).

### 3.2 Single-contrast vs. multi-task localizers: Empirical example

To extend our findings from simulation to real data, we used the example of functional localization of the language network in the right cerebellar hemisphere. In a recent paper, Casto et al. (2026) used the single-contrast of sentence reading vs. non-word reading to localize this region. First, we replicated their results using our language multi-task battery dataset, which included the sentence reading and non-word reading tasks. To quantify the relationship between estimated ROI size and fSNR, we first computed the sentence *>* non-words contrast based on 8 min of data for each subject (see 2.8). We then used a range of t-value thresholds to assign voxels of the right cerebellar hemisphere to the language region based on this contrast. We also estimated the cerebellar fSNRs for each subject, using all data and tasks, except the two tasks used in the localizer (see 2.5). Consistent with our simulations, the average correlation between fSNR and ROI size was *r* = 0.59, *p* = 0.01, averaged across all tested thresholds (see 2.8 for details). This result confirmed that individuals with better data quality had larger estimated language ROIs.

For the multi-task localizer, we selected five tasks from the language multi-task battery dataset: sentence reading, theory of mind, tongue movement, demand grid and action observation (action). We used activity estimates for these tasks to match the overall length of the single-contrast localizer (8 minutes per subject). The tasks were selected because they have been previously reported to activate the cerebellar language regions and neighboring areas (Nettekoven et al., 2024). We then estimated individual parcellations by assigning all voxels of the right cerebellar hemisphere either to the language ROI or four other competing ROIs (see 2.8). Unlike the single-contrast localizer, the correlation between fSNR and the size of the multi-task language ROI was not significant (*r* = − 0.34, *p* = 0.19), indicating that region size estimates from the multi-task approach are not biased by data quality.

To compare the different localizers directly, we calibrated the thresholds for the single-contrast localizer so that the average estimated ROI size matched the average estimated ROI size for the multi-task localizer. This yielded a fixed t-threshold of 1.47, and an adaptive threshold of the top 10.4% voxels in each individual.

We then assessed how consistently the cerebellar language region was identified across individuals, by calculating the percentage of individuals in which each voxel was assigned to the language ROI. All three localizers defined ROIs that included the medial part of Crus I and II. However, the overlap between individuals was stronger for the multi-task (Fig. 3b) than for the single-contrast localizers (Fig. 3a). To evaluate this, we computed the average Dice coefficient of each individual ROI mask with the ROI masks of each other individual (Fig. 3c). The average Dice coefficient was significantly higher for the multi-task localizer (average Dice = 0.26±0.01) than for the contrast-fixed (average Dice = 0.12 ±0.01, *t*_16_ = 13.83, *p* = 2.57e-10) and contrast-adaptive localizer (average Dice = 0.12 ±0.003, *t*_16_ = 15.45, *p* = 4.92e-11). Thus, for comparable region size and amount of individual imaging data, the multi-task localizer led to a spatially more consistent selection of the cerebellar language region in right medial crus I and II (denoted language III in Casto et al. (2026), and region S1R and S2R in Nettekoven et al. (2024)).

**Figure 3.**
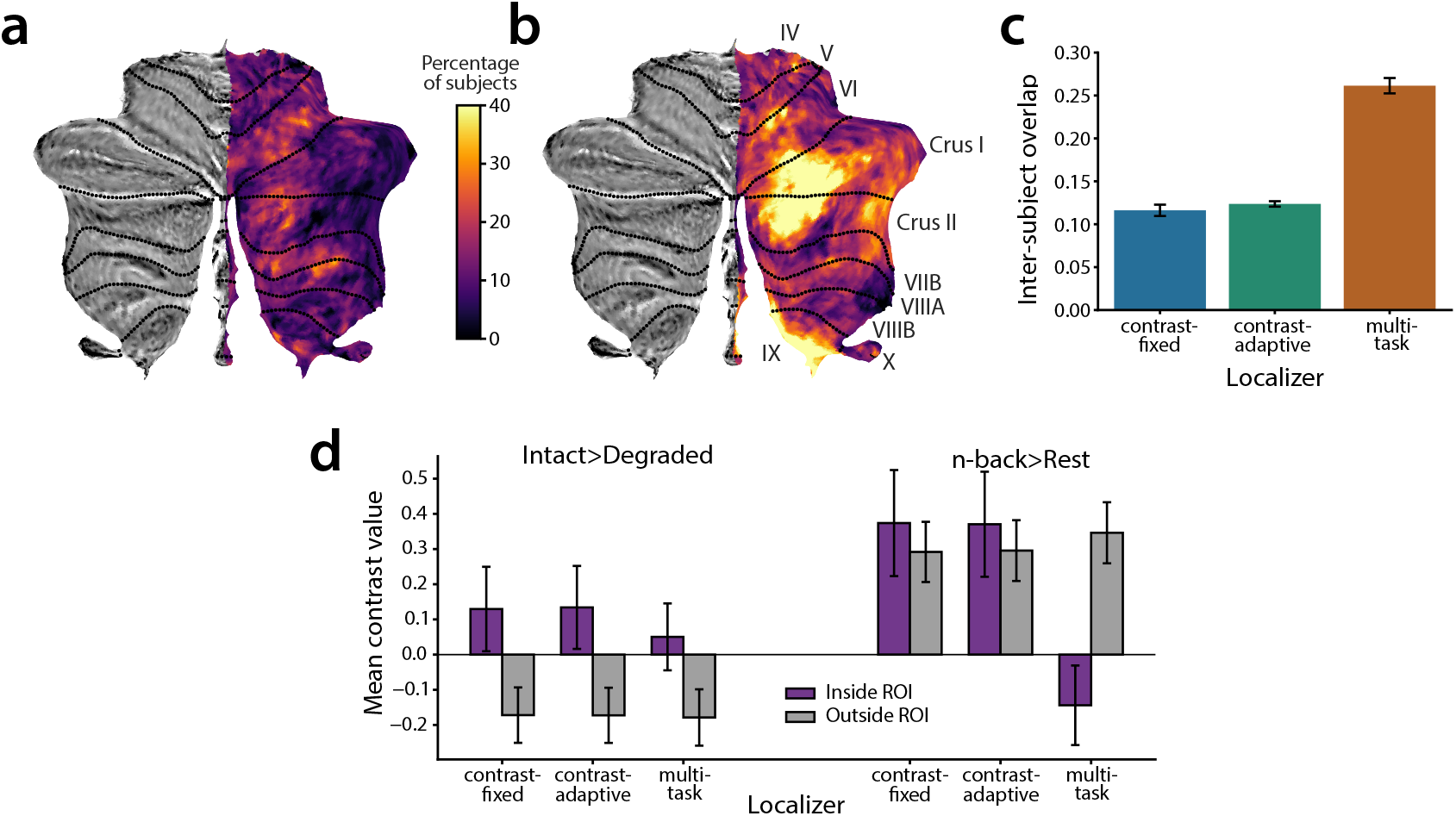
Empirical validation in cerebellar language localization. **(a)** Percentage of individuals in which each cerebellar voxel was assigned to the language ROI using the contrast-fixed localizer. **(b)** Percentage of individuals in which each cerebellar voxel was assigned to the language ROI using a multi-task localizer. **(c)** Average inter-subject Dice coefficient for each localizer. Error bars indicate standard error of the mean across subjects. **(d)** Selectivity and specificity of single-contrast and multi-task language localizers in the cerebellum. Mean activation across individuals for a target and a control contrast inside and outside the defined language ROI (constrained to the right cerebellar hemisphere). Error bars indicate standard error of the mean across subjects.

The largest difference between the two localizers was that the single-contrast localizers not only selected voxels in medial Crus I and II, but also selected voxels in lobule VI, VIIb and VIIIa. These regions are also activated by the language-specific contrast, but were also strongly engaged during working memory and executive function tasks, and are therefore considered part of the multi-demand network (Nettekoven et al., 2024).

To understand the functional differences between the selected ROIs, we looked at the mean activation for two types of contrasts that were not included in either the single-contrast or multi-task localizer. First, we compared the activity for listening to intact passages vs. degraded passages. Like the single-contrast localizer, this represents a language-comprehension contrast (Casto et al., 2026). A good language ROI should therefore also reveal a positive contrast. Indeed, we found that voxels inside of the putative language ROI showed the expected higher activity for intact passages compared to degraded passages, independent of the localizer method. The contrast did not show any difference between the multi-task and contrast-fixed localizers, *t*_16_ = −0.95, *p* = 0.36, and between the multi-task and contrast-adaptive localizers, *t*_16_ = −1.02, *p* = 0.32 (Fig. 3d).

We then asked how specifically the selected ROI was activated by language functions. Because neighboring areas were also activated by working memory tasks, we selected the activity of the n-back task vs. rest as our control contrast. On this task, the multi-task localizer showed substantially lower activity when compared to the single-contrast localizers for the control contrast inside the putative language ROI (purple solid bars) *t*_16_ = −4.87, *p* = 1.71e-4 for multi-task vs contrast-fixed and *t*_16_ = −4.66, *p* = 2.63e-4 for multi-task vs contrast-adaptive.

Thus, the single-contrast localizer selected voxels that showed high activity on the language-specific contrast, but also often show high activity for working memory tasks. The multi-task localizer excluded those voxels due to the inclusion of a working memory task (Demand grid) in the battery and the competitive approach to region assignment. This led to a spatially more consistent and functionally more specific ROI definition. While both approaches can be further improved by taking into account spatial information (see discussion), based on functional data alone, the multi-task localizer led to functionally more consistent ROI definition. Additionally, it also addressed the problem of the threshold and fSNR dependency of the single-contrast localizer.

### 3.3 Optimal battery selection: Simulation

While multi-task batteries can be advantageous even when localizing a single functional ROI, their main strength becomes apparent when one wants to parcellate a brain structure into multiple functional ROIs or model connectivity between two brain structures. In our companion paper, we show that the multi-task approach consistently outperforms the use of an equivalent amount of resting-state data for such applications (Nettekoven et al., 2026).

When designing a multi-task battery for a specific application, however, the researcher must address two critical questions: how many tasks should be included, and which specific tasks should be chosen? In many cases, these questions will be constrained by practical considerations. Some tasks cannot be included, because they are not feasible for the study population. Other tasks have to be included to allow comparisons to external datasets. However, in many cases the researcher will have some flexibility in battery design. Here we propose the idea that, if we know the average activity patterns elicited by each task in the brain structure of interest, we can use a data-driven approach to assemble the optimal task battery to characterize that structure.

Imagine you had a large task library of 100 possible tasks, each of which had a known activity profile across 5 functionally distinct regions. How would we best choose the set of tasks that would optimally detect the boundaries between these regions in single subjects? To address this question, we compared three different battery selection strategies to assemble hypothetical task batteries containing anywhere between *B* = 3−16 different tasks.

As a baseline, we used a *Random* strategy, simply selecting the *B* tasks without replacement at random from the 100 tasks. For the *Maximal Contrast* method, we picked the combination of tasks that resulted in the largest activations relative to rest across tasks, averaged across all regions. However, to distinguish functional regions, we do not only want to maximize the overall activation, we also want to minimize the similarities across regions. One way to achieve this is to think about the response profile for each region as a regressor in a linear model, and then to minimize the collinearity between these regressors (*Minimal Collinearity*, see section 2.10 for details). We then tested the different selection strategies in two application scenarios: Parcellation and connectivity modeling.

For parcellation, we asked how well we could functionally parcellate the brain structure of interest into functional regions using data acquired with this task battery. For each task battery, we generated noisy fMRI data (assuming 8 min of scan time) for a hypothetical brain structure containing the 5 functional regions. Using many simulated datasets and the known activity profile for each region, we then estimated a parcellation for each battery and compared it to the ground truth parcellation using the Dice overlap.

First, we found that data-driven battery selection strategy clearly outperformed random battery selection (Fig. 4a). Second, as the number of unique tasks increased, the quality of the estimated parcellation improved, but this relationship plateaued after about 8 tasks. Third, the criterion that maximized the differences between the tasks (*Minimal Collinearity*) slightly outperformed the criterion that maximized contrasts between task and rest (*Maximal Contrast*). Finally, it appeared that the difference between the data-driven selection strategies converged with an increasing number of tasks.

**Figure 4.**
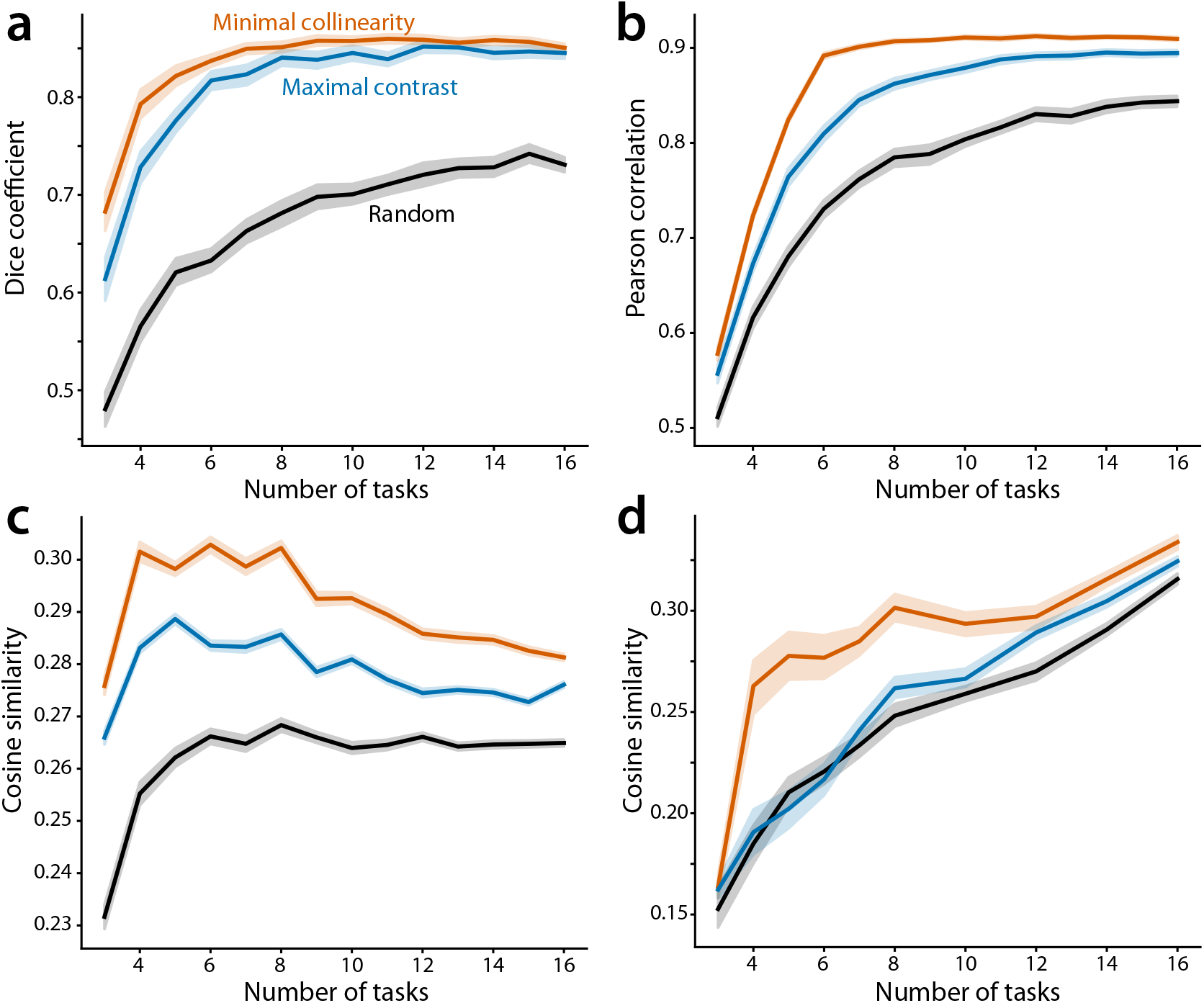
Optimal battery selection strategies across battery sizes. **(a)** Parcellation simulations. Dice overlap between estimated and true parcellations. Shaded area indicates standard deviation across 100 simulations with different task libraries. **(b)** Connectivity modeling simulations. Correlation between true connectivity weights and estimated connectivity weights. Shaded area indicates standard deviation across 100 simulations with different task libraries. **(c)** Parcellation of the neocortex in fMRI. Cosine similarity between true test data and predicted test data, averaged across subjects. Shaded area indicates standard error across subjects (N=24) of the differences between selection strategies in the MDTB dataset. **(d)** Neocortex-cerebellum connectivity modeling in fMRI. Cosine similarity between true cerebellar test data and predicted cerebellar test data, averaged across subjects. Shaded area indicates standard error across subjects of the differences between selection strategies in the MDTB dataset.

We then replicated these results for a connectivity model. Here, the goal was to estimate the directed, effective connectivity weights from a source structure to a target structure. In this type of model we want to account for the influence of the different source regions onto a single target region, such that we can predict the activity of the target region from the activity of a source region (Cole et al., 2016; Reid et al., 2019). Here, we evaluated performance by correlating the recovered connectivity weights using a specific battery with the ground-truth connectivity weights.

The results follow a similar pattern as in the parcellation simulations. Data-driven battery selection strategies outperformed random battery selection, and *Minimal Collinearity* outperformed *Maximal Contrast* (Fig. 4b). Performance improved with an increasing number of tasks but started to plateau at around 8 tasks.

### 3.4 Optimal battery selection: Empirical examples

To test whether the advantages seen in the simulations hold under real conditions, we applied the same battery selection framework to the MDTB dataset. We used task set B of the MDTB to construct different task batteries of 8 min length each, and used these data to estimate individual parcellations and individual connectivity models. We then evaluated these on task set A. Importantly, both the estimation of these brain mapping models and the evaluation was done within each individual.

For parcellation, we attempted to subdivide the neocortex of each individual into functionally distinct regions. We used the 360 parcels of the Glasser atlas (Glasser et al., 2016) to define neocortical functional profiles (see section 2.12). We then estimated functional parcellations for each individual using only the data for the selected task battery, and evaluated how well this parcellation could predict the data on a different task set for the same subject. Performance was measured as the average cosine similarity of the true and predicted test data across vertices and subjects.

As in our simulations, parcellations based on data-driven battery selection strategies generally outperformed random selection after averaging performance across battery sizes (*t*_23_ = 10.54, *p* = 2.78e-10 for *Maximal Contrast* vs *Random, t*_23_ = 13.86, *p* = 1.18e-12 for *Minimal Collinearity* vs *Random*) (Fig. 4c). Second, maximizing the differences between the regions (*Minimal Collinearity*) outperformed *Maximal Contrast* (*t*_23_ = 9.36, *p* = 2.62e-9). Finally, performance reached its peak at around 5-7 tasks - after that adding more unique tasks slightly reduced performance. We replicated these results when parcellating only parts of the neocortex (e.g., prefrontal cortex) or subcortical structures such as the cerebellum (Supplementary fig. S2).

For connectivity modeling, we built a directed connectivity model from 1002 neocortical regions to each voxel of the cerebellum (King et al., 2023). We estimated ridge regression connectivity models for each individual using the candidate task batteries based on task set B. The resulting connectivity models were used to predict the cerebellar data in task set A, using only the neocortical data acquired on the same tasks (see section 2.12). Performance was measured as the cosine similarity between the true and predicted cerebellar test data.

Again, data-driven battery selection outperformed random selection after averaging performance across battery sizes (Fig. 4d), (*t*_23_ = 8.14, *p* = 3.20e-8 for *Minimal Collinearity* vs *Random*). For this scenario, using task batteries that maximized the functional contrast, however, only marginally outperformed random battery selection, *t*_23_ = 2.08, *p* = 0.05. Maximizing the differences between regions (*Minimal Collinearity*) outperformed *Maximal Contrast* after averaging across battery sizes (*t*_23_ = 6.94, *p* = 4.47e-7). In contrast to the parcellations, for which performance peaked at 5-7 tasks, connectivity models continued to improve performance as the number of unique tasks increased.

The advantage of data-driven selection strategies over random selection was greatest in the 4-8 task range for both parcellations and connectivity models (Fig. 4c-d). To test whether the data-driven selection advantage was battery-size dependent, we compared *Minimal Collinearity* to *Random* selection using a two-way repeated measures ANOVA with selection type and battery size as the interaction factors. We found a strong selection type x battery size interaction effect for parcellations (*F*_13,299_ = 53.20, *p <* 0.001) and for connectivity modeling (*F*_13,299_ = 24.81, *p <* 0.001).

In summary, we found that the best data-driven strategy for battery selection is to maximize the size of the contrasts between tasks and the mean activity, while at the same time minimizing the similarities between the different regions within the brain structure. Of course, the optimal set of tasks will depend both on the available tasks in the task library, and on the part of the brain that we wish to characterize. With the release of the multi-task toolbox, we also publish a large task library with 56 task conditions. As an example, we applied our minimal collinearity approach to build batteries of different sizes for the sensori-motor areas, the prefrontal cortex, and the cerebellum (see 2.9 and supplementary table S1).

### 3.5 Optimal design of Multi-task battery experiments: Grouped vs. interspersed designs

Once a task battery is chosen, the next question is how to best design an fMRI experiment to measure activity for these tasks. One approach is to *group* similar tasks into specific runs or sessions, as was done in the HCP (Barch et al., 2013) and IBC dataset (Pinho et al., 2018). This design has the advantage that participants can concentrate on groups of similar tasks (for example motor localizers) within each imaging run. It has, however, the drawback that most task-to-task comparisons have to be made between, rather than within, imaging runs. As we will show here, such an approach can be statistically suboptimal.

The alternative design is therefore to randomly *intersperse* all tasks, with each task condition appearing once within each imaging run (King et al., 2019). While this design demands from the participant a constant switching between tasks, which can be somewhat demanding, we will show that it has much better statistical characteristics for task-to-task comparison.

The advantage of the interspersed fMRI design stems from the fact that we cannot measure fMRI activity signal in absolute terms, but only as a change relative to a baseline state, usually phases of rest. When we want to estimate task-related activation, we have to contend with two sources of measurement noise: The variability due to the measurement of the task itself 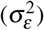, and the variability involved in measuring the baseline activity 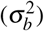. As the baseline is usually common to all tasks tested within a single run, measurement noise on the baseline also introduces a positive covariance between all tasks measured in a single run. This positive covariance can be clearly seen in different sessions of the HCP task dataset (Fig. 5a). By using the different elements of the covariance matrix, we can quantify the importance of the measurement noise of the task and the measurement noise of the baseline (see 2.13). Across the different sessions of the HCP task dataset, the baseline measurement accounted for between 22.1% (Working Memory) and 86.7% (Language) of the total measurement noise of the activity estimates (Fig. 5b).

**Figure 5.**
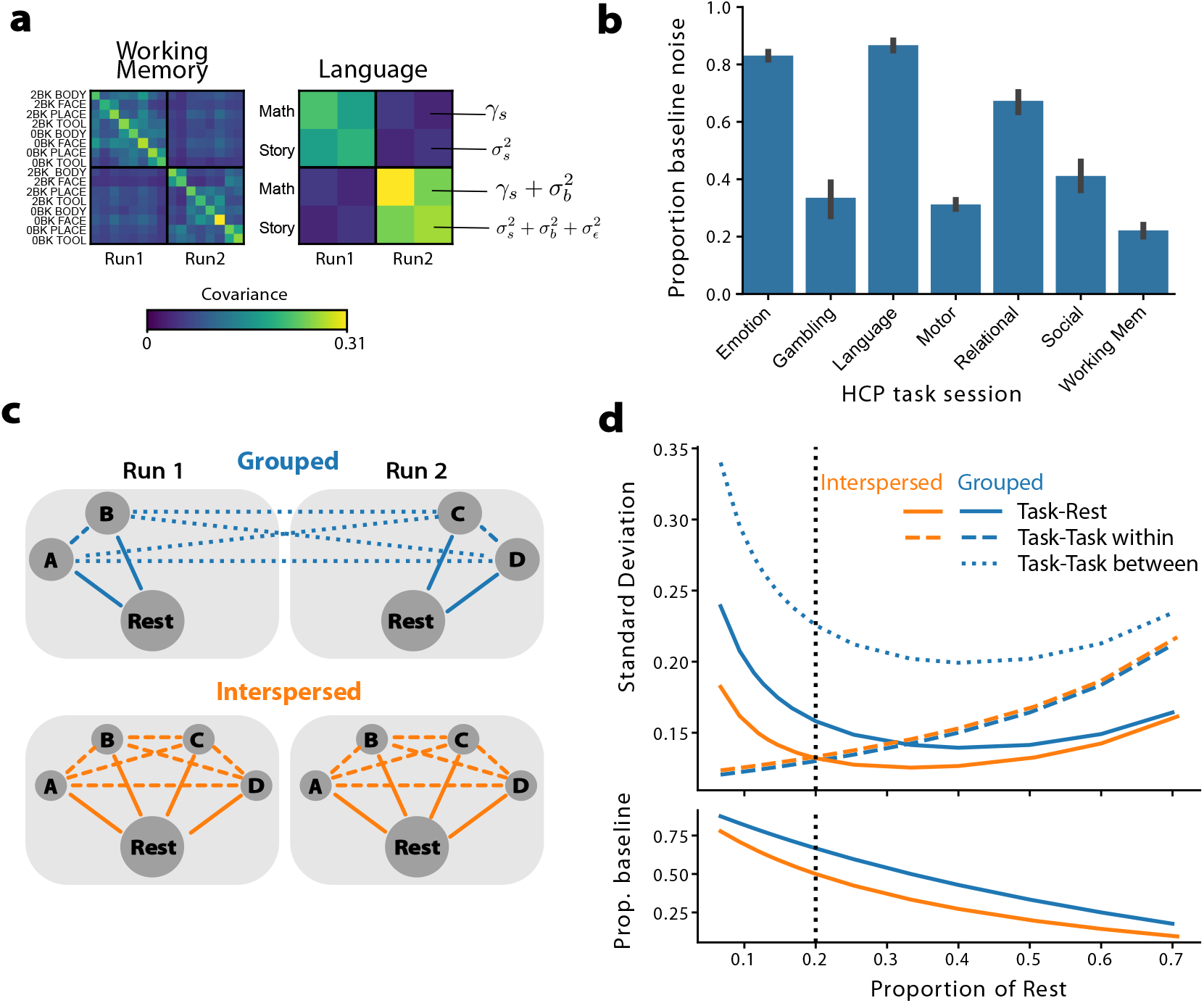
Experimental design of multi-task batteries. **(a)** Average covariance of neocortical activity patterns across two imaging runs for the working memory and the language sessions of the HCP task dataset. **(b)** Proportion of the estimated baseline noise variance 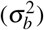relative to the total estimated noise variance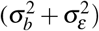across the different sessions of the HCP task dataset. Error bars indicate SEM across participants. **(c)** In a grouped design, only some task-task contrasts can be made within-run (dashed lines), while most contrasts have to be made between runs (dotted line). In an interspersed design, all contrasts can be made within-run. **(d)** Predicted standard deviation of task-rest, and task-task contrasts for grouped (blue) and interspersed (orange) designs as a function of the proportion of each imaging run dedicated to rest. Lower panel shows the proportion of noise variance that is due to estimation of resting baseline. For the vertical line, the length of rest is equivalent to each single task condition.

How does the baseline variance influence the variability of our contrasts of interest? To obtain an analytical insight, we simulated a scenario with 2 imaging runs and 4 tasks. In the grouped design (Fig. 5c), two tasks were scanned in run 1, and the other two tasks in run 2. In the interspersed design, both runs included all the tasks. The total length of each run was kept constant across the two designs. We also varied the proportion of time that was spent on measuring the resting baseline. We adjusted the simulation parameters, such that the baseline measurement noise accounted for, depending on the amount of rest, 17.6% 81.8% of the variance of the activity estimates, thereby covering the range spanned by different HCP task sessions (Fig. 5d).

Because the two designs allocated the same amount of time to each task (either within a single run or across two runs), the within-run task-task contrasts (Fig. 5d, dashed line) are measured with the same variability in both designs. As resting baseline is not important here, the lowest variability would be achieved in a design that does not include rest at all. In contrast, the between-run task-task contrasts in the grouped design (dotted line) rely on the common baseline measurement across the two runs. As it is impacted by the measurement noise on the baseline for both run 1 and 2, its variance is more than twice as high as compared to the within-run contrast. In the interspersed design all contrasts can be performed within-run, making the high-variance between-run comparisons unnecessary.

Maybe surprisingly, the interspersed design also results in less variable estimates for the (within-run) contrasts between task and rest (solid line). This is due to the fact that in the grouped design, the contrast can only be calculated against the resting baseline of one run, while the interspersed design can leverage the resting baseline in both runs.

Finally, we show that the advantage of the interspersed design remains when we take into account the time lost by switching between tasks within runs (Supplementary fig. S3).

### 3.6 Temporal autocorrelation in interspersed designs

In the above analysis, we used the average covariance across tasks for imaging runs with a length of 2:16-5:01 min to estimate the variance induced by the measurement of a common baseline. However, larger task batteries require longer imaging runs. For example, King et al. (2019) used 17 tasks per task set, necessitating imaging runs of 10 min length. This length allows for 35s for each task block, a good design choice given the low-pass filtering properties of the hemodynamic response Friston et al. (1999). However, long imaging runs have the problem that low-frequency drift in the fMRI signal (Lund et al., 2006) will add to the variability of contrasts between tasks that are temporally very far separated within a run.

To analyze this problem, we used the MDTB dataset to estimate the covariance of the measurement noise between different tasks within the same run, depending on how far apart they were temporally spaced (see 2.15). The higher this covariance is, the larger will be the benefit of using a within-run vs. a between-run contrast. As can be seen from Fig. 6a, the measurement covariance was highest for tasks that were immediately neighboring, and then fell off to reach an asymptote for a separation of approximately 7 tasks (4 min). Nonetheless, the measurement covariance even for large temporal separations did not reach zero, reflecting the fact that all tasks within each run are still measured with respect to the same common baseline (rest).

**Figure 6.**
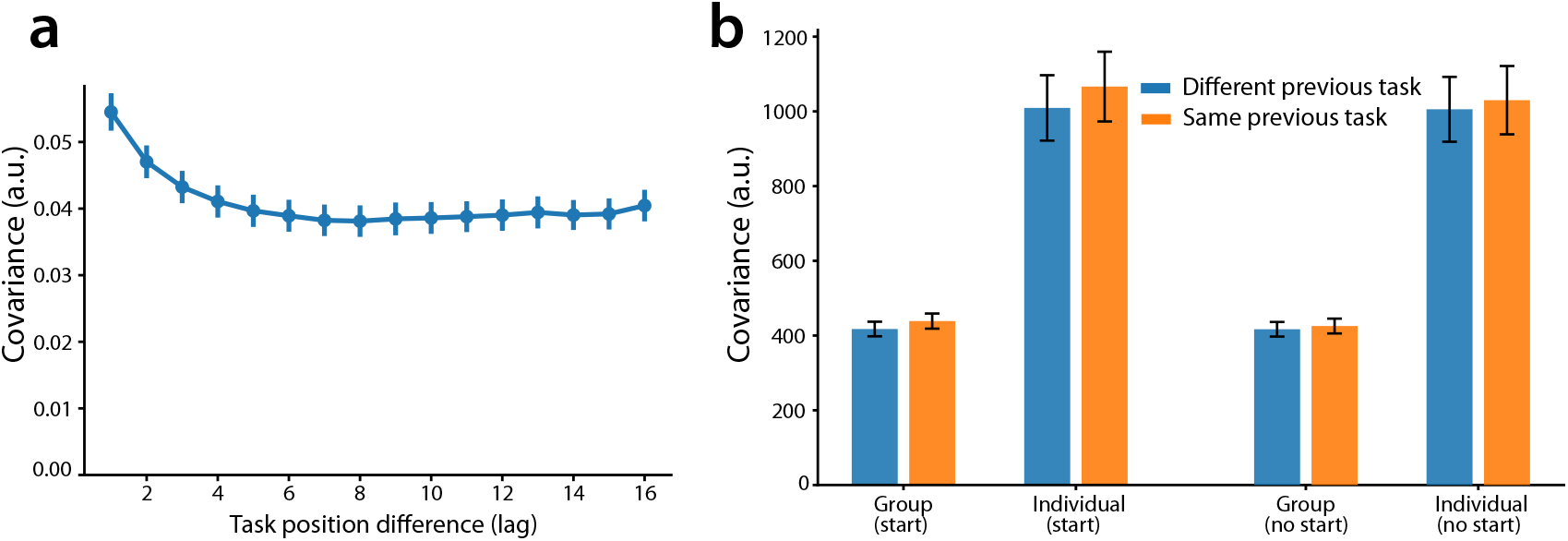
Temporal autocorrelation and task carry-over effects in interspersed designs. **(a)** Mean covariance of task-evoked activity patterns in the MDTB dataset as a function of the difference in task positions (lag) within each imaging run. Error bars indicate standard error of the mean across subjects. **(b)** Mean covariance of the task-evoked activity patterns when a task is preceded by a different task vs the same task, shown for both across subjects (group) and within subject (individual) estimates. Error bars indicate standard error of the mean across subjects.

This analysis showed that the advantage of interspersed designs over grouped designs will diminish somewhat when imaging runs exceed 4 min of length. However, even in this case, a substantial part of the measurement variability will be due to the measurement of the common baseline. Thus, the maximal length of the imaging run, and hence the maximal size of a task battery, can mostly be determined by practical considerations.

### 3.7 Task carry-over in interspersed designs

Another concern when combining many different tasks into a single run is carry-over effects, i.e. that the estimate of task activity for a task may change systematically depending on which task came before it. To measure the size of such effects, we again used the MDTB dataset, in which 17 tasks were scanned in 16 runs in different order. We calculated covariance between the measured activity patterns for the same task across different runs, depending on whether it was preceded by the same or different task (Fig. 6b). We conducted this analysis within the same individuals, highlighting any subject-specific carry-over effect, and also between different individuals, highlighting carry-over effects that are systematic across the entire group. We found that the estimated covariance between neocortical task estimates was significantly larger when the tasks were preceded by the same than different tasks (*t*_23_ = 7.97, *p* = 4.57e-8 for group and *t*_23_ = 3.37, *p* = 0.003 for individual), indicating the presence of significant order or carry-over effects. However, compared to the systematic variance in the data explained by the tasks, the amount of variance explained by the previous task was only 4.9± 0.5% for group and 5.0 ±1.5% for individual.

Some of these effects, however, could be due to the fact that the first task of each run had especially high or low activity. We therefore repeated the analysis, this time removing the first task in each run. In the group analysis the estimated covariance again was smaller when the task was preceded by a different task (*t*_23_ = 3.33, *p* = 0.003), but this difference was not significant anymore within individuals (*t*_23_ = 1.48, *p* = 0.15). The amount of variance explained by the previous task compared to the total systematic variance was 2.0± 0.6% at the group level.

Thus, these results indicate that there are small, but reliable carry-over effects between tasks in a run. About half of this carry-over effect can be attributed to the first task in the imaging run. Because full counter-balancing of the sequence across many tasks is not practical, we recommend randomizing the order of tasks across subjects. Overall, however, the carry-over effects are small relative to the systematic task-related variance, such that even subjects acquired with different task sequences can be confidently compared.

## 4 DISCUSSION

In this paper, we advocate the use of multi-task batteries for individual brain mapping. Our companion paper showed that multi-task fMRI outperforms resting-state fMRI in estimating parcellations or connectivity models that can predict new brain states in the same individuals (Nettekoven et al., 2026). Here, we compare the multi-task approach against the commonly adopted single-contrast functional localizer (Kanwisher et al., 1997; Fedorenko et al., 2010; Dodell-Feder et al., 2011; Scott et al., 2017). We show that, even when localizing a single region, multi-task localizers produce functional ROIs that are spatially more consistent across individuals. Multi-task batteries have the additional advantage of being threshold independent, and can localize multiple functional regions at the same time. To help the design of good task batteries, we derive, develop and validate data-driven strategies for task selection. Finally, we address important experimental design questions for implementing multi-task batteries.

### Multi-task vs. single-contrast localizers

One core advantage of the multi-task over the single-contrast localizer approach is that it identifies functional regions by detecting the likely boundaries to neighboring regions. This is essential if one wishes to compare the size of functional regions across individuals. For example, it has been shown that the size of early visual regions can differ more than threefold across individuals (Benson et al., 2022). When using single-contrast localizers the estimated size of the region is highly dependent on the statistical threshold: When using the same threshold for each individual, individuals with higher functional fSNR will have more voxels assigned to a functional region, leading to a bias in the estimated region size. When using an adaptive threshold, one can enforce the same region size across individuals, but this approach is insensitive to true size differences. The multi-task approach is able to identify regions with different sizes, as it assigns voxels based on the direction of the response profile rather than the magnitude, thereby competitively choosing between the target region and neighboring regions, and hence detecting the most likely position of the functional boundary.

Another difference between the two localizer approaches lies in the nature of the regions they identify. Single-contrast localizers highlight the full network of areas that are engaged by a given cognitive function. In the example used in this paper -identifying the language regions in the human cerebellum-this leads to the identification of 4 spatially separate areas that are also functionally heterogeneous (Casto et al., 2026). Three of these areas show strong activity for executive function tasks, with only one (LangCereb3 in Crus I/II/lobule VIIb) being selectively activated by language material. In contrast, the multi-task approach identifies this functionally homogeneous parcel based on the response profile of these voxels, and by contrasting it with other regions that show high activity during executive function tasks. Thus, if the goal is to identify the complete network of regions activated by a function, single-contrast localizers remain appropriate; but if the goal is to isolate a specific, functionally coherent region, multi-task batteries are preferable.

Both the single-contrast and the multi-task approach can be improved by combining individual subject data with group-level spatial information. In the single-contrast approach, Fedorenko and colleagues used a Group-Constrained Subject-Specific (GSS) approach (Fedorenko et al., 2010; Julian et al., 2012), in which the average group contrast map is used to define spatially segregated regions. This hard group parcellation is then used to subdivide individual localizer maps, thereby constraining individual ROIs to be fully enclosed in a specific spatial region. The multi-task approach has been combined with a hierarchical Bayesian framework that uses a probabilistic group parcellation as a spatial prior that is then combined with the individual data (the conditional) to compute an integrated estimate (Zhi et al., 2025). Both approaches bias the individual parcellations towards the group map and will therefore improve the spatial overlap of the selected ROI across individuals. However, the probabilistic approach will still work when there is no clear and hard spatial separation between two functional regions, for example when dissociating two components of the default-mode network (Braga and Buckner, 2017), which are highly overlapping on the group level. The Bayesian approach (Zhi et al., 2025) explicitly quantifies the uncertainty of the anatomical information, and can optimally integrate functional and anatomical information.

### Design considerations

The remainder of our results provides new insights into the optimal design of multi-task batteries. Assuming that the overall scan time per subject is limited, we show that including more tasks generally leads to parcellation and connectivity models that generalize better to other sets of tasks (see also Nettekoven et al., 2026). However, the specific choice of tasks also matters: Optimally, each task in the battery activates the target structure in a unique way. We show that a mathematical criterion that minimizes the collinearity (*Minimal Collinearity*) provides a selection criterion that improves the ability of both parcellations and connectivity models to predict novel activity patterns in novel sets of tasks. This directly enables the construction of optimal multi-task batteries using a data driven, empirical approach. Of course, in many cases the exact choice of tasks is dictated by theoretical or practical considerations.

For example, in most cases researchers will want to include both rest (as a standard fMRI baseline) and one or more tasks of interest. However, even in such cases, there is usually some flexibility in choosing the remaining tasks to complete the battery. Here a data-driven criterion can help compare different possible designs, thereby helping to pick the best companion tasks for the battery of interest.

Our results also show that it is optimal to intersperse all tasks within a single run (King et al., 2019). This approach improves the reliability of most task-to-task contrasts, as they can now be computed directly within each run, rather than between runs. There are of course some drawbacks with the interspersed design: First, it is slightly more demanding for participants, as it requires continuous switching between cognitively distinct tasks. This can lead to increased fatigue and may not be suitable for all populations. Second, each task transition requires an instruction period to remind the participants of the upcoming tasks, which slightly reduces the total time available for measuring the activity during task performance. Third, interspersed designs require the researchers to commit to a full task battery before data collection begins, preventing a cumulative approach where researchers can decide to add additional tasks in later sessions without having to run all previous tasks (Pinho et al., 2018).

Despite these practical drawbacks, we believe that the interspersed design of multi-task battery localizers offers such a powerful statistical advantage that it is worth the extra effort in design and participant instruction.

### Implementation and open software tools

To ease the implementation of this approach, we are releasing the open-source python toolbox Multi- TaskBattery (Arafat et al., 2026). Built upon PsychoPy Peirce (2007); Peirce et al. (2019), the toolbox implements optimal battery selection, stimulus presentation, response collection, recording of behavioral and eye-tracking data, scanner synchronization, and instructions. With 20+ tasks currently implemented, the toolbox allows for the flexible and fast assembly of new batteries. With an object-oriented design, the toolbox can be easily extended with new tasks and response devices.

The multi-task battery approach can be especially powerful when multiple datasets spanning various functional domains can be integrated. To ease such analysis, we established a data management framework, entitled FunctionalFusion, that utilizes BIDS-derivative standards to bring different multi-task battery or deep phenotyping datasets into a common analysis framework. Currently, we have 10 multi-task datasets, including the Multi-domain task battery King et al. (2019), the individual brain charting project Pinho et al. (2018), the Human Connectome task dataset Barch et al. (2013), and the 103 task dataset Nakai and Nishimoto (2020), processed in the framework. The python package enables the quick extraction of functional contrasts and time series in any desired group atlas space. Using this toolbox, our group has already published a paper that integrates results from 111 individuals and 417 different task conditions into a functional atlas of the human cerebellum (Nettekoven et al., 2024).

### Best practices when designing multi-task batteries

In summary, our results suggest the following best practices for designing and analyzing multi-task fMRI batteries:

- If the goal of the study is to identify connectivity models or derive a comprehensive functional parcellation of an individual, multi-task designs are preferable to resting-state fMRI (Nettekoven et al., 2026).
- If the goal of the study is to identify a single homogeneous functional region, multi-task designs are preferable to traditional functional localizers, especially if the size of the individual functional regions is a variable of interest.
- Task batteries with a larger number of tasks tend to generalize better to new sets of tasks. Based on our results, a battery size of 7 or more tasks already allows for stable generalization performance.
- When selecting tasks, we recommend choosing tasks that activate the structure of interest in as many different ways as possible. Using a library of tasks and associated activation patterns, we recommend using a data driven criterion (*Minimal Collinearity*) to optimize battery selection.
- Including rest as a task is not necessary but possible depending on experimental needs. Here, we rely on the contrast between tasks, so rest can just be treated as another task.
- For practical reasons we recommend the inclusion of *anchor* tasks in the multi-task battery (for example rest) that allow the comparison to other datasets. After choosing those tasks, the remaining tasks can be optimized using a data-driven approach.
- As much as possible, we recommend using interspersed designs where all tasks are performed within each run. This optimizes the contrast between all task pairs and removes the impact of baseline measurement noise.
- The sequence of tasks should be randomized across runs to avoid introducing carry-over effects between tasks. Overall, however, these effects are small relative to the task-specific signal (2-5 % of task variance).
- Because the interspersing of tasks within a run can be cognitively demanding, we recommend familiarizing the subjects before the scanning session with all the tasks that require a response, and to conclude with at least 2 training runs, in which all tasks are presented in a random sequence.

### Practical applications

Multi-task fMRI batteries offer a scalable approach to characterizing brain function across multiple functional domains within a single scanning session. While not exhaustive, we highlight two ways in which this framework may prove particularly useful: (1) Presurgical functional mapping (2) Brain-behavior modeling.

Although fMRI presurgical mapping has traditionally relied on rsfMRI due to its ease of acquisition and minimal cognitive demand on the patient, current advances in task-based mapping make it increasingly practical and certainly more informative than rsfMRI for presurgical planning. The framework introduced in this paper allows seamless task selection and paradigm design depending on the specific brain structure of interest. This means that a surgeon can define the structure (or part of a structure) of interest and get a set of tasks tailored to their specific problem. An additional consideration for presurgical mapping is the amount of time the patient needs to be in the MRI scanner in order to acquire data that is reliable enough. Recent work showed that combining even a limited amount of individual data (10 minutes) with a probabilistic group parcellation using Bayesian approaches produces robust individualized maps (Zhi et al., 2025; Nettekoven et al., 2024). These advances lower the barrier for adoption of multi-task batteries in the clinical setting.

A second application area where multi-task batteries have strong potential is modeling brain-behavior relationships. Traditionally, studies of brain-behavior relationships have been limited by sample size and amount of scanning time used per participant, making it difficult to detect reliable brain-behavior associations. Recent work has highlighted that many of the findings in the literature are based on statistically underpowered paradigms, usually with a small sample size (Varoquaux, 2018; Marek et al., 2022). The multi-task framework addresses this by encouraging the use of task batteries that in many cases share tasks across studies and datasets. As more research groups adopt this approach and share the resulting task contrasts in a common analysis framework, it becomes feasible to aggregate datasets at scale. The inclusion of a number of standardized “anchor” tasks across many of these datasets is particularly valuable, as it allows researchers to assemble large diverse data structures covering many individuals across many task conditions. This can ultimately enable well-powered studies of how brain activation patterns relate to cognitive traits, psychiatric disorders and clinical outcomes.

### Limitations and future directions

In this paper, we proposed a comprehensive framework for studying the individual organization of the brain using multi-task fMRI. Here, we validated this framework using data collected on healthy young adults. Although the general framework is broadly applicable, the extension of our approach to clinical populations is just now beginning. Some of the tasks included in the library are currently only well suited to healthy young adults, but may be too cognitively demanding for children, older adults or clinical populations. To implement the framework for clinical studies, many of the tasks included in the task library can be adjusted parametrically, allowing researchers to carefully adjust the difficulty level of each task based on the behavioral performance of the population of interest.

Finally, even though we have implemented a range of tasks, the current library still only covers a small proportion of the tasks and activities humans perform in everyday life. Continued expansion of this library through collaboration and data sharing will be necessary to enable data-driven strategies to decide on truly representative batteries and to more comprehensively address the problem of task-dependency of the mapping results.

## 5 DATA AVAILABILITY

The raw fMRI data of the Language task battery dataset has been deposited in the OpenNeuro database under accession code ds007276 (DOI:10.18112/openneuro.ds007276.v1.1.0). The raw fMRI data of the MDTB dataset is available under accession code ds002105 (DOI:10.18112/openneuro.ds002105.v1.1.0). For the HCP dataset, raw data is available at https://www.humanconnectome.org/. The pro-cessed fMRI activity patterns are available at Zenodo for the Language task battery https://doi.org/10.5281/zenodo.19100385, MDTB https://doi.org/10.5281/zenodo.16788784, and HCP https://doi.org/10.5281/zenodo.16903949 datasets. The combined task library is available at Zenodo https://doi.org/10.5281/zenodo.18793343.

## 6 CODE AVAILABILITY

The open source software toolbox for designing and running multi-task batteries (including stimulus presentation and response collection) is available at https://github.com/DiedrichsenLab/MultiTaskBattery.git. The framework for managing and integrating a range of multiple multi-task fMRI datasets is available at https://github.com/DiedrichsenLab/Functional_Fusion. The specific analysis code for generating the results and figures in this paper is available at https://github.com/DiedrichsenLab/OptimalBattery.git.

## ACKNOWLEDGMENTS

This work was supported by a Discovery Grant from the Natural Sciences and Engineering Research Council of Canada (NSERC, RGPIN-2016-04890), and a project grant from the Canadian Institutes of Health Research (CIHR, PJT-191815), both to J.D. Additional funding came from the Canada First Research Excellence Fund (BrainsCAN) to Western University and the Raynor Cerebellum Project.

C.N. was supported by a Wellcome Trust Early Career Award (306553/Z/23/Z) and a Junior Research Fellowship Grant from Linacre College, University of Oxford.

## AUTHOR CONTRIBUTIONS

B.A., C.N. and J.D. conceived the study, C.N. and J.D. guided the study design, B.A. collected the data, J.D. guided the analysis, J.X. provided data analysis tools, B.A. performed the analysis, B.A. and J.D. wrote the manuscript, all authors reviewed and approved the manuscript.

## 7 SUPPLEMENTARY MATERIALS

### 7.1 Group vs individual covariance matrices

To derive battery selection metrics, we estimate a task-by-task covariance matrix from the fMRI activity patterns. In the main analyses in this paper, we estimated the covariance matrix using an individual-level cross-validated approach: for each individual in the MDTB dataset, we computed the covariance matrix by multiplying the activity patterns from different runs only, and then averaged the resulting matrices across individuals.

This approach differs from a simpler group-averaged approach in two ways. First, it captures individual differences between the tasks that may only be visible at the individual subject level, but are obscured at the group level because functionally distinct regions can be spatially tightly interdigitated (Braga and Buckner, 2017; Xue et al., 2021). Second, cross-validation ensures that the measurement noise does not inflate the diagonal elements of the covariance matrix, providing more accurate task contrast estimates.

In practice, however, combining tasks from two different datasets requires using group-averaged activity patterns from each dataset, since individuals differ across datasets, and within-subject covariances cannot be computed across datasets. Here, we tested how much such a group-based estimate of the covariance matrix differs from the more precise individual-level cross-validated covariance matrix.

Using neocortical data in fs32k space from the MDTB dataset session B, we compared the two approaches. First, we calculated the cross-validated covariance matrix of each individual and averaged them to get a single covariance matrix (Fig. S1a). Second, we computed the covariance matrix directly from the group-averaged activity patterns (Fig. S1b). The two matrices were highly correlated (*r* = 0.99, *p <* 0.001), indicating that the group-based covariance matrix captures the covariance structure present at the individual level. This allows researchers to build libraries of task activity patterns measured in separate groups of individuals without requiring access to individual data or run-wise data.

### 7.2 Open task library

To facilitate the adoption of multi-task batteries, we provide a publicly available library containing group-averaged activity patterns for 56 task conditions estimated from a total of 41 individuals (available at https://doi.org/10.5281/zenodo.18793343). This library combines tasks from two datasets used in this paper: the Language Task Battery and the Multi-Domain Task Battery (MDTB).

Combining group-averaged activity patterns across datasets with different individuals requires ad-dressing two main points. First, all datasets must share a common baseline. Here, all activity patterns were expressed relative to the resting baseline before merging. Second, differences in signal scaling can arise from differences in scanning parameters and pre-processing pipelines. We address this by requiring each dataset to share at least one “anchor” task with another dataset. Starting with a root dataset (here, MDTB) as the initial library, each new dataset is calibrated by computing a scaling factor that minimizes the squared difference between all the shared conditions in the new dataset and their equivalents in the current library. This factor is applied to all activity patterns of the new dataset before merging. Shared conditions are then combined using a weighted average based on the number of individuals in each dataset contributing to the final activity estimate, while unique conditions are added directly. This procedure will allow us to extend the library with new datasets, as long as rest is included as a baseline and they include at least one anchor task that is common with the existing library.

**Figure S1.**
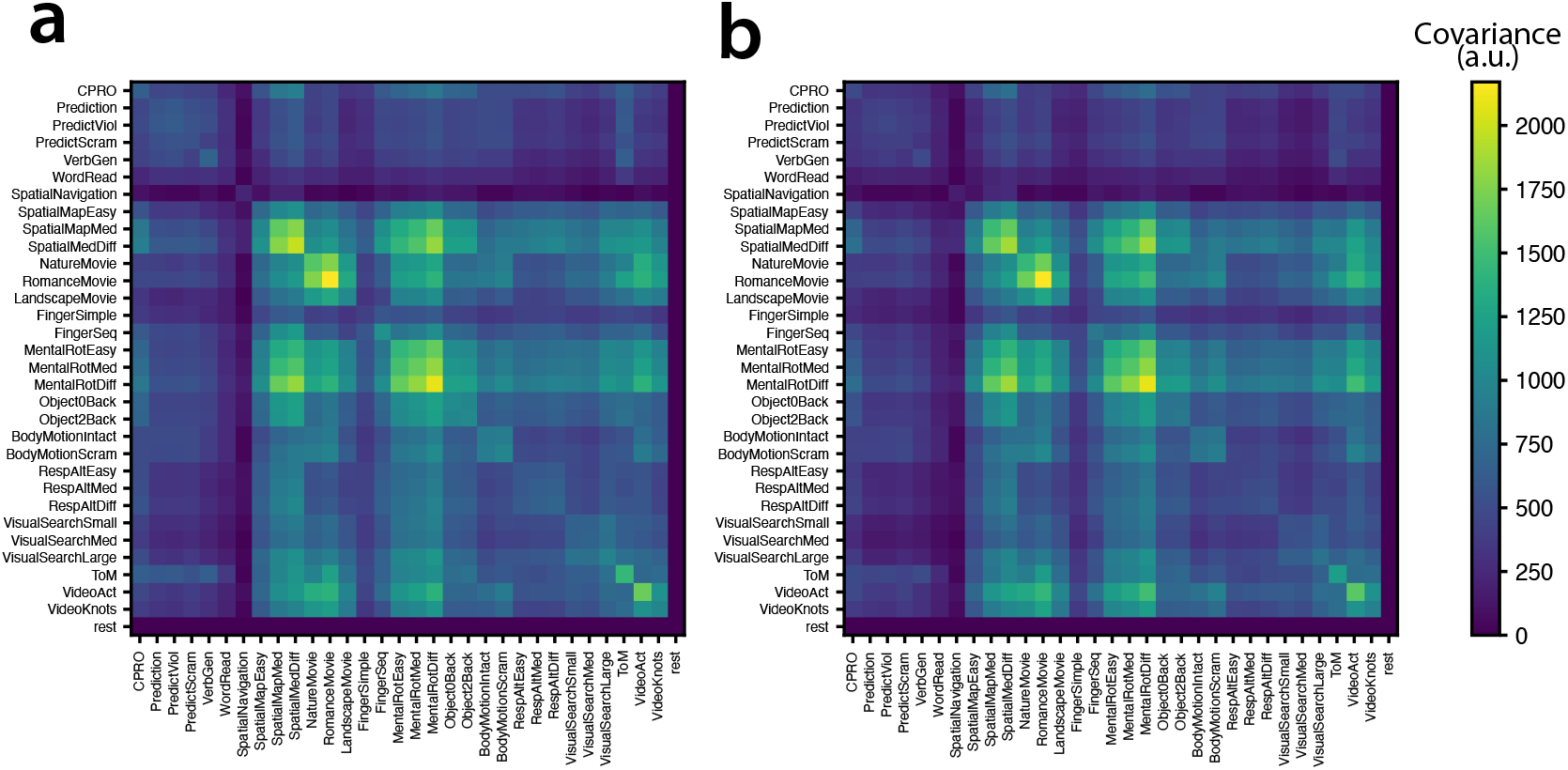
Group vs. individual task-by-task covariance matrices. **(a)** Covariance matrix using an individual-level cross-validated approach. **(b)** Covariance matrix using group-averaged data without cross-validation. This library can be directly used in the MultiTaskBattery toolbox to select optimal task batteries for a brain structure of interest. Researchers can specify the target brain structure and the number of tasks, and the toolbox will apply the *Minimal Collinearity* criterion described in this paper to recommend tasks suitable for brain mapping studies.

The library of task activity patterns is shared following the Human Connectome Project conventions. Neocortical activity patterns are provided as vertices in fs32k space and subcortical structures - including cerebellum, hippocampus, amygdala, thalamus, brainstem and basal ganglia - are provided as voxels in MNI152NLin6Asym space at 2mm resolution.

**Supplementary table S1.** Example optimal task batteries by region and battery size for an 8-minute fMRI scan.

| Battery size | Motor | PFC | Cerebellum |
| --- | --- | --- | --- |
| 4 tasks | finger sequence, tongue movement, action observation (action), rest | verb generation, romance movie, mental rotation (difficult), rest | finger sequence, tongue movement, emotional face (happy), rest |
| 6 tasks | finger sequence, tongue movement, action observation (action), go no go (go), mental rotation (medium), rest | verb generation, theory of mind, 2-back pic, 2-back verbal, biomotion normal, rest | finger sequence, tongue movement, theory of mind, 2-back verbal, emotional face (sad), rest |
| 8 tasks | finger sequence, tongue movement, action observation (action), go no go (go), motor imagery, emotional face (happy), 2-back pic, rest | verb generation, theory of mind, 2-back pic, 2-back verbal, affective (pleasant), go no go (go), biomotion normal, rest | verb generation, action observation, finger sequence, tongue movement, 2-back verb, mental rotation (medium), emotional face (happy), rest |

**Figure S2.**
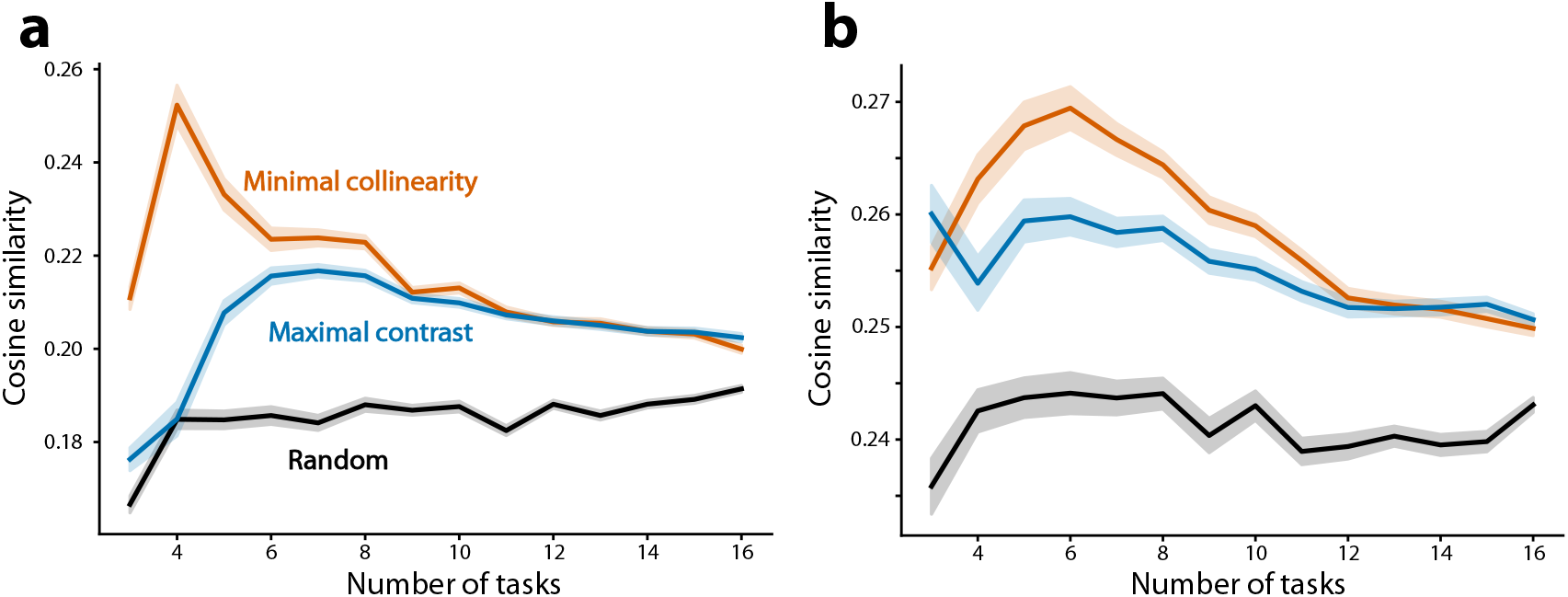
Optimal battery selection strategies for brain parcellations of different brain structures. **(a)** Prefrontal cortex. Cosine similarity between predicted and measured test data, averaged across subjects. Shaded area indicates standard error across subjects of the differences between selection strategies in the MDTB dataset. Regions are defined using 44 parcels of the Glasser atlas (Glasser et al., 2016; Donahue et al., 2018). **(b)** Cerebellum. Regions defined using the NettekovenSym32 atlas (Nettekoven et al., 2024).

**Figure S3.**
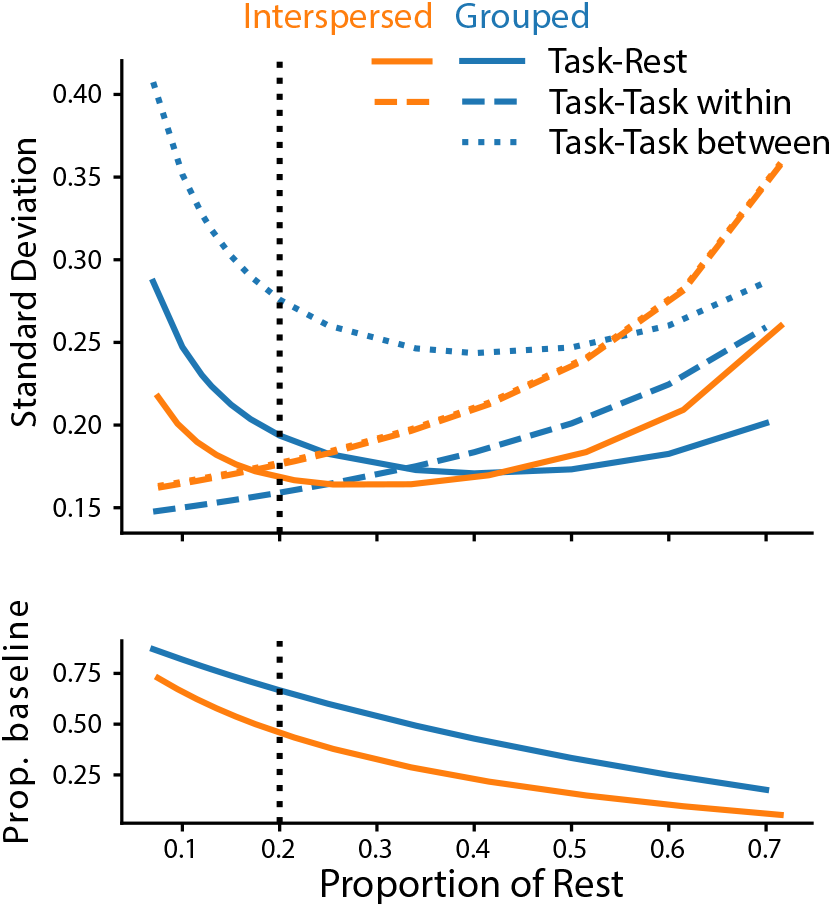
Grouped vs Interspersed designs after accounting for instruction periods. Predicted standard deviation of task-rest, and task-task contrasts for grouped (blue) and interspersed (orange) designs as a function of the proportion of each imaging run dedicated to rest. Lower panel shows the proportion of noise variance that is due to estimation of resting baseline. For the vertical line, the length of rest is equivalent to each single task condition.

**Supplementary table S2.** Language task battery descriptions and feedback details.

| **Task Name** | **Task Description** | **Feedback** |
| --- | --- | --- |
| Rest | Lie in the scanner looking at a fixation cross | NA |
| Action observation | Watch videos of knot tying (Cross et al., 2012). Conditions:<br>(1) Action: Observe a knot being tied<br>(2) Knot: Passively view the knot in a 360-degree-rotating view | NA |
| Verb Generation | Silently read visually presented nouns. Conditions:<br>(1) Read: Passively read the noun<br>(2) Generate: Covertly generate an associated verb | NA |
| Intact Passage | Listen to audible sentences (Scott et al., 2017) | NA |
| Degraded Passage | Listen to muffled sentences (Scott et al., 2017) | NA |
| Sentence Reading | Read sentences word by word (Fedorenko et al., 2010) | NA |
| Non-words Reading | Read non-words (Fedorenko et al., 2010) | NA |
| Auditory Narrative | Listen to a narrative (Moth Radio Hour) | NA |
| Theory of Mind | Read 2-sentence stories about other people, answer short questions | Trial + task |
| Tongue Movement | Move tongue alternating from right to left at 1Hz (Saadon-Grosman et al., 2022) | NA |
| Finger Sequence | Perform a randomly generated six-element finger sequence. | Trial + task |
| Romance Movie | Watch clips of the movie “Up”, without audio or subtitles | NA |
| N-back | Judge if the current stimulus is the same as the one presented 2 serial positions back | Trial + task |
| Oddball | Identify red ‘K’ from red or black ‘K’ or ‘O’ (Du et al., 2024) | Task |
| Demand Grid | Solve complex spatial grid (Fedorenko et al., 2013) | Trial + task |
| Spatial Navigation | Imagine navigating in a childhood home from one location to another | NA |

## REFERENCES

Arafat, B., Nettekoven, C., and Diedrichsen, J. (2026). Multitaskbattery: A python toolbox for multi-task fmri battery design.

Barch, D. M., Burgess, G. C., Harms, M. P., Petersen, S. E., Schlaggar, B. L., Corbetta, M., Glasser, M. F., Curtiss, S., Dixit, S., Feldt, C., Nolan, D., Bryant, E., Hartley, T., Footer, O., Bjork, J. M., Poldrack, R., Smith, S., Johansen-Berg, H., Snyder, A. Z., and Van Essen, D. C. (2013). Function in the human connectome: task-fMRI and individual differences in behavior. Neuroimage, 80:169–189.

Benson, N. C., Yoon, J. M. D., Forenzo, D., Engel, S. A., Kay, K. N., and Winawer, J. (2022). Variability of the surface area of the V1, V2, and V3 maps in a large sample of human observers. J. Neurosci., 42(46):8629–8646.

Biswal, B. B. and Uddin, L. Q. (2025). The history and future of resting-state functional magnetic resonance imaging. Nature, 641(8065):1121–1131.

Braga, R. M. and Buckner, R. L. (2017). Parallel interdigitated distributed networks within the individual estimated by intrinsic functional connectivity. Neuron, 95(2):457–471.e5.

Buckner, R. L., Krienen, F. M., and Yeo, B. T. T. (2013). Opportunities and limitations of intrinsic functional connectivity MRI. Nat. Neurosci., 16(7):832–837.

Casto, C., Poliak, M., Tuckute, G., Small, H., Sherlock, P., Wolna, A., Lipkin, B., D’Mello, A. M., and Fedorenko, E. (2026). The cerebellar components of the human language network. Neuron.

Chen, G., Cai, Z., Kording, K. P., Liu, T., Faskowitz, J., Bandettini, P. A., Biswal, B., and Taylor, P. A. (2025). Resting-state fMRI and the risk of overinterpretation: Noise, mechanisms, and a missing rosetta stone. bioRxiv.

Cole, M. W., Ito, T., Bassett, D. S., and Schultz, D. H. (2016). Activity flow over resting-state networks shapes cognitive task activations. Nat. Neurosci., 19(12):1718–1726.

Cross, E. S., Cohen, N. R., Hamilton, A. F. d. C., Ramsey, R., Wolford, G., and Grafton, S. T. (2012). Physical experience leads to enhanced object perception in parietal cortex: insights from knot tying. Neuropsychologia, 50(14):3207–3217.

Dice, L. R. (1945). Measures of the amount of ecologic association between species. Ecology, 26(3):297– 302.

Diedrichsen, J. (2006). A spatially unbiased atlas template of the human cerebellum. Neuroimage, 33(1):127–138.

Dodell-Feder, D., Koster-Hale, J., Bedny, M., and Saxe, R. (2011). fMRI item analysis in a theory of mind task. Neuroimage, 55(2):705–712.

Donahue, C. J., Glasser, M. F., Preuss, T. M., Rilling, J. K., and Van Essen, D. C. (2018). Quantitative assessment of prefrontal cortex in humans relative to nonhuman primates. Proceedings of the National Academy of Sciences, 115(22):E5183–E5192.

Du, J., DiNicola, L. M., Angeli, P. A., Saadon-Grosman, N., Sun, W., Kaiser, S., Ladopoulou, J., Xue, A., Yeo, B. T. T., Eldaief, M. C., and Buckner, R. L. (2024). Organization of the human cerebral cortex estimated within individuals: networks, global topography, and function. J. Neurophysiol., 131(6):1014–1082.

Fedorenko, E., Duncan, J., and Kanwisher, N. (2013). Broad domain generality in focal regions of frontal and parietal cortex. Proc. Natl. Acad. Sci. U. S. A., 110(41):16616–16621.

Fedorenko, E., Hsieh, P.-J., Nieto-Castañón, A., Whitfield-Gabrieli, S., and Kanwisher, N. (2010). New method for fMRI investigations of language: defining ROIs functionally in individual subjects. J. Neurophysiol., 104(2):1177–1194.

Fischl, B. (2012). FreeSurfer. Neuroimage, 62(2):774–781.

Friston, K. J., Zarahn, E., Josephs, O., Henson, R. N., and Dale, A. M. (1999). Stochastic designs in event-related fMRI. Neuroimage, 10(5):607–619.

Glasser, M. F., Coalson, T. S., Robinson, E. C., Hacker, C. D., Harwell, J., Yacoub, E., Ugurbil, K., Andersson, J., Beckmann, C. F., Jenkinson, M., Smith, S. M., and Van Essen, D. C. (2016). A multi-modal parcellation of human cerebral cortex. Nature, 536(7615):171–178.

Glasser, M. F., Sotiropoulos, S. N., Wilson, J. A., Coalson, T. S., Fischl, B., Andersson, J. L., Xu, J., Jbabdi, S., Webster, M., Polimeni, J. R., Van Essen, D. C., Jenkinson, M., and WU-Minn HCP Consortium (2013). The minimal preprocessing pipelines for the human connectome project. Neuroimage, 80:105– 124.

Gordon, E. M., Laumann, T. O., Adeyemo, B., Huckins, J. F., Kelley, W. M., and Petersen, S. E. (2016). Generation and evaluation of a cortical area parcellation from resting-state correlations. Cereb. Cortex, 26(1):288–303.

Gordon, E. M., Laumann, T. O., Gilmore, A. W., Newbold, D. J., Greene, D. J., Berg, J. J., Ortega, M., Hoyt-Drazen, C., Gratton, C., Sun, H., Hampton, J. M., Coalson, R. S., Nguyen, A. L., McDermott,K. B., Shimony, J. S., Snyder, A. Z., Schlaggar, B. L., Petersen, S. E., Nelson, S. M., and Dosenbach, N. U. F. (2017). Precision functional mapping of individual human brains. Neuron, 95(4):791–807.e7.

Gratton, C., Laumann, T. O., Nielsen, A. N., Greene, D. J., Gordon, E. M., Gilmore, A. W., Nelson, S. M., Coalson, R. S., Snyder, A. Z., Schlaggar, B. L., Dosenbach, N. U. F., and Petersen, S. E. (2018). Functional brain networks are dominated by stable group and individual factors, not cognitive or daily variation. Neuron, 98(2):439–452.e5.

Julian, J. B., Fedorenko, E., Webster, J., and Kanwisher, N. (2012). An algorithmic method for functionally defining regions of interest in the ventral visual pathway. Neuroimage, 60(4):2357–2364.

Kanwisher, N., McDermott, J., and Chun, M. M. (1997). The fusiform face area: a module in human extrastriate cortex specialized for face perception. J. Neurosci., 17(11):4302–4311.

King, M., Hernandez-Castillo, C. R., Poldrack, R. A., Ivry, R. B., and Diedrichsen, J. (2019). Functional boundaries in the human cerebellum revealed by a multi-domain task battery. Nat. Neurosci., 22(8):1371– 1378.

King, M., Shahshahani, L., Ivry, R. B., and Diedrichsen, J. (2023). A task-general connectivity model reveals variation in convergence of cortical inputs to functional regions of the cerebellum. Elife, 12.

Kong, R., Li, J., Orban, C., Sabuncu, M. R., Liu, H., Schaefer, A., Sun, N., Zuo, X.-N., Holmes, A. J.,Eickhoff, S. B., and Yeo, B. T. T. (2019). Spatial topography of individual-specific cortical networks predicts human cognition, personality, and emotion. Cereb. Cortex, 29(6):2533–2551.

Kong, R., Yang, Q., Gordon, E., Xue, A., Yan, X., Orban, C., Zuo, X.-N., Spreng, N., Ge, T., Holmes, A., Eickhoff, S., and Yeo, B. T. T. (2021). Individual-specific areal-level parcellations improve functional connectivity prediction of behavior. Cereb. Cortex, 31(10):4477–4500.

Laumann, T. O., Gordon, E. M., Adeyemo, B., Snyder, A. Z., Joo, S. J., Chen, M.-Y., Gilmore, A. W., McDermott, K. B., Nelson, S. M., Dosenbach, N. U. F., Schlaggar, B. L., Mumford, J. A., Poldrack, R. A., and Petersen, S. E. (2015). Functional system and areal organization of a highly sampled individual human brain. Neuron, 87(3):657–670.

Lund, T. E., Madsen, K. H., Sidaros, K., Luo, W.-L., and Nichols, T. E. (2006). Non-white noise in fMRI: does modelling have an impact? Neuroimage, 29(1):54–66.

Marek, S., Tervo-Clemmens, B., Calabro, F. J., Montez, D. F., Kay, B. P., Hatoum, A. S., Donohue, M. R., Foran, W., Miller, R. L., Hendrickson, T. J., Malone, S. M., Kandala, S., Feczko, E., Miranda-Dominguez, O., Graham, A. M., Earl, E. A., Perrone, A. J., Cordova, M., Doyle, O., Moore, L. A., Conan, G. M., Uriarte, J., Snider, K., Lynch, B. J., Wilgenbusch, J. C., Pengo, T., Tam, A., Chen, J., Newbold, D. J., Zheng, A., Seider, N. A., Van, A. N., Metoki, A., Chauvin, R. J., Laumann, T. O., Greene, D. J., Petersen, S. E., Garavan, H., Thompson, W. K., Nichols, T. E., Yeo, B. T. T., Barch,D. M., Luna, B., Fair, D. A., and Dosenbach, N. U. F. (2022). Reproducible brain-wide association studies require thousands of individuals. Nature, 603(7902):654–660.

Mueller, S., Wang, D., Fox, M. D., Yeo, B. T. T., Sepulcre, J., Sabuncu, M. R., Shafee, R., Lu, J., and Liu, H. (2013). Individual variability in functional connectivity architecture of the human brain. Neuron, 77(3):586–595.

Nakai, T. and Nishimoto, S. (2020). Quantitative models reveal the organization of diverse cognitive functions in the brain. Nat. Commun., 11(1):1142.

Nettekoven, C., Shahbazi, A., Arafat, B., Skenderija, M., Xiang, J. D., Luisa Pinho, A., and Diedrichsen, J. (2026). Multi-task fMRI outperforms resting-state fMRI for revealing task-invariant organization of the human brain. bioRxiv.

Nettekoven, C., Zhi, D., Shahshahani, L., Pinho, A. L., Saadon-Grosman, N., Buckner, R. L., and Diedrichsen, J. (2024). A hierarchical atlas of the human cerebellum for functional precision mapping. Nat. Commun., 15(1):8376.

Peirce, J., Gray, J. R., Simpson, S., MacAskill, M., Höchenberger, R., Sogo, H., Kastman, E., and Lindeøv, J. K. (2019). PsychoPy2: Experiments in behavior made easy. Behav. Res. Methods, 51(1):195–203.

Peirce, J. W. (2007). PsychoPy–psychophysics software in python. J. Neurosci. Methods, 162(1-2):8–13.

Pinho, A. L., Amadon, A., Ruest, T., Fabre, M., Dohmatob, E., Denghien, I., Ginisty, C., Becuwe-Desmidt,S., Roger, S., Laurier, L., Joly-Testault, V., Médiouni-Cloarec, G., Doublé, C., Martins, B., Pinel, P., Eger, E., Varoquaux, G., Pallier, C., Dehaene, S., Hertz-Pannier, L., and Thirion, B. (2018). Individual brain charting, a high-resolution fMRI dataset for cognitive mapping. Scientific data, 5.

Reid, A. T., Headley, D. B., Mill, R. D., Sanchez-Romero, R., Uddin, L. Q., Marinazzo, D., Lurie, D. J., Valdés-Sosa, P. A., Hanson, S. J., Biswal, B. B., Calhoun, V., Poldrack, R. A., and Cole, M. W. (2019). Advancing functional connectivity research from association to causation. Nat. Neurosci., 22(11):1751–1760.

Saadon-Grosman, N., Angeli, P. A., DiNicola, L. M., and Buckner, R. L. (2022). A third somatomotor representation in the human cerebellum. bioRxiv.

Schaefer, A., Kong, R., Gordon, E. M., Laumann, T. O., Zuo, X.-N., Holmes, A. J., Eickhoff, S. B., and Yeo, B. T. T. (2018). Local-global parcellation of the human cerebral cortex from intrinsic functional connectivity MRI. Cereb. Cortex, 28(9):3095–3114.

Scott, T. L., Gallée, J., and Fedorenko, E. (2017). A new fun and robust version of an fMRI localizer for the frontotemporal language system. Cogn. Neurosci., 8(3):167–176.

Varoquaux, G. (2018). Cross-validation failure: Small sample sizes lead to large error bars. Neuroimage, 180(Pt A):68–77.

Wang, D., Buckner, R. L., Fox, M. D., Holt, D. J., Holmes, A. J., Stoecklein, S., Langs, G., Pan, R., Qian, T., Li, K., Baker, J. T., Stufflebeam, S. M., Wang, K., Wang, X., Hong, B., and Liu, H. (2015). Parcellating cortical functional networks in individuals. Nat. Neurosci., 18(12):1853–1860.

Worsley, K. J. and Friston, K. J. (1995). Analysis of fMRI time-series revisited–again. Neuroimage, 2(3):173–181.

Xue, A., Kong, R., Yang, Q., Eldaief, M. C., Angeli, P. A., DiNicola, L. M., Braga, R. M., Buckner, R. L., and Yeo, B. T. T. (2021). The detailed organization of the human cerebellum estimated by intrinsic functional connectivity within the individual. J. Neurophysiol., 125(2):358–384.

Yeo, B. T. T., Krienen, F. M., Sepulcre, J., Sabuncu, M. R., Lashkari, D., Hollinshead, M., Roffman, J. L., Smoller, J. W., Zöllei, L., Polimeni, J. R., Fischl, B., Liu, H., and Buckner, R. L. (2011). The organization of the human cerebral cortex estimated by intrinsic functional connectivity. J. Neurophysiol., 106(3):1125–1165.

Zhi, D., Shahshahani, L., Nettekoven, C., Pinho, A. L., Bzdok, D., and Diedrichsen, J. (2025). A hierarchical bayesian brain parcellation framework for fusion of functional imaging datasets. Imaging Neuroscience, 3.

